# Inherited *Brain Fitness* Deficit caused by Light Stress Promotes Glioblastoma Progression in Drosophila

**DOI:** 10.1101/2025.08.28.672790

**Authors:** Teresa de los Reyes Corrales, Clara E. Gavira-O’Neill, Marta Portela, Esther Seco, Carlos Rodríguez-Martín, Sergio Casas-Tintó

## Abstract

Environmental and lifestyle factors can influence long-term human health outcomes, including the development of complex diseases such as cancer. Glioblastoma is the most aggressive and fatal form of brain cancer, marked by its invasive nature and resistance to standard therapies. Despite similar genetic alterations, patients diagnosed with glioblastoma often show highly variable survival outcomes, suggesting that non-genetic influences may play a critical role in tumor progression.

Here we show that environmental conditions experienced by the parental line, such as permanent exposure to artificial light, can affect the brain’s resilience to tumor growth in the next generation. This effect, described as reduced *Brain fitness*, does not impair brain development but accelerates glioma progression and shortens survival in the offspring.

These findings point to a novel concept, the *Brain fitness* hypothesis, in which inherited susceptibility to brain tumors may be shaped by parental exposures. Our results suggest that non-coding RNAs could mediate this intergenerational transmission, offering new insight into the complexity of inherited traits beyond direct genetic mutations. This work broadens current understanding of glioma biology and highlights the importance of environmental health across generations.

**Abstract:** Environmental exposure including modern lifestyle habits and occupational exposures describe a wide spectrum of risk factors for human health including carcinogenesis. Glioblastoma is the most common, aggressive, and lethal type of glioma. It is highly proliferative and invasive, infiltrates the surrounding brain parenchyma and is resistant to current treatments. Patients suffering from glioblastoma have a wide heterogeneity in life expectancy, even with similar mutations, that can vary from weeks to years. Many potential risk factors for glioma have been studied to date, but few provide significant correlations for tumor prognosis. Here, we provide a novel perspective on the interaction between the tumor and host depending on paternal environmental or lifestyle determinants, coined as *Brain fitness hypothesis*. Permanent light exposure induces genetic stress hallmarks in the progeny that do not alter central nervous system development, but compromise *Brain fitness* to an accelerated glioma growth, ultimately reducing life expectancy. Reduced survival in the progeny correlates with an increase in the number of activated brain genes involved in neuronal dynamics as well as those relevant for glioma progression such as presynaptic, circadian genes or genes required for the tumoral network. Among the inheritable mechanisms, ncRNAs emerge as potential vectors for the transmission of parental exposure to the subsequent generations and might explain the complexity of inheritable traits and phenotypes acquired as a pleiotropic effect.

BOX OF DEFINITIONS
**Light pollutome/exposome:** The sum of environmental exposures or modern lifestyle habits as a source of artificial lighting or solar radiation that compromise fitness and disease progression from early stages (prenatal/postnatal) to adulthood.
**Luminic/Light stress (LL):** Continuous light exposure (24h) at low luminance intensity (1,000 lux) of white light.
***Brain fitness hypothesis*:** Molecular factors that precondition brain susceptibility to pathological aggressions modulated by environmental or lifestyle determinants.
**Luminic stress effects (LSE):** Genetic changes in unexposed F1 brains in response to pre-conceptional parental luminic exposure as a memory of parental experiences.
**Intergenerational Inheritance: Transmission of** parental(F0) environmental effects to the descendants (F1or F2 depending on gestational state).

**Graphical abstract:** Maternal pre-cexceptional exposure to light stress (1000 lux white light) induces intergenerational changes in the offspring brain fitness leading on a sex dependent light stress memory hallmark (LSE). Thus, F1 flies acquire a brain with the fitness tatus altered and preconditioned to GB development Consequently, GB grows faster and causes worse prognosis in affected animals. In this context miRNAs emerge as a potential epigenetic mechanism mediating this intergenerational effect, describing rriRNA-979 expression as a risk factor.

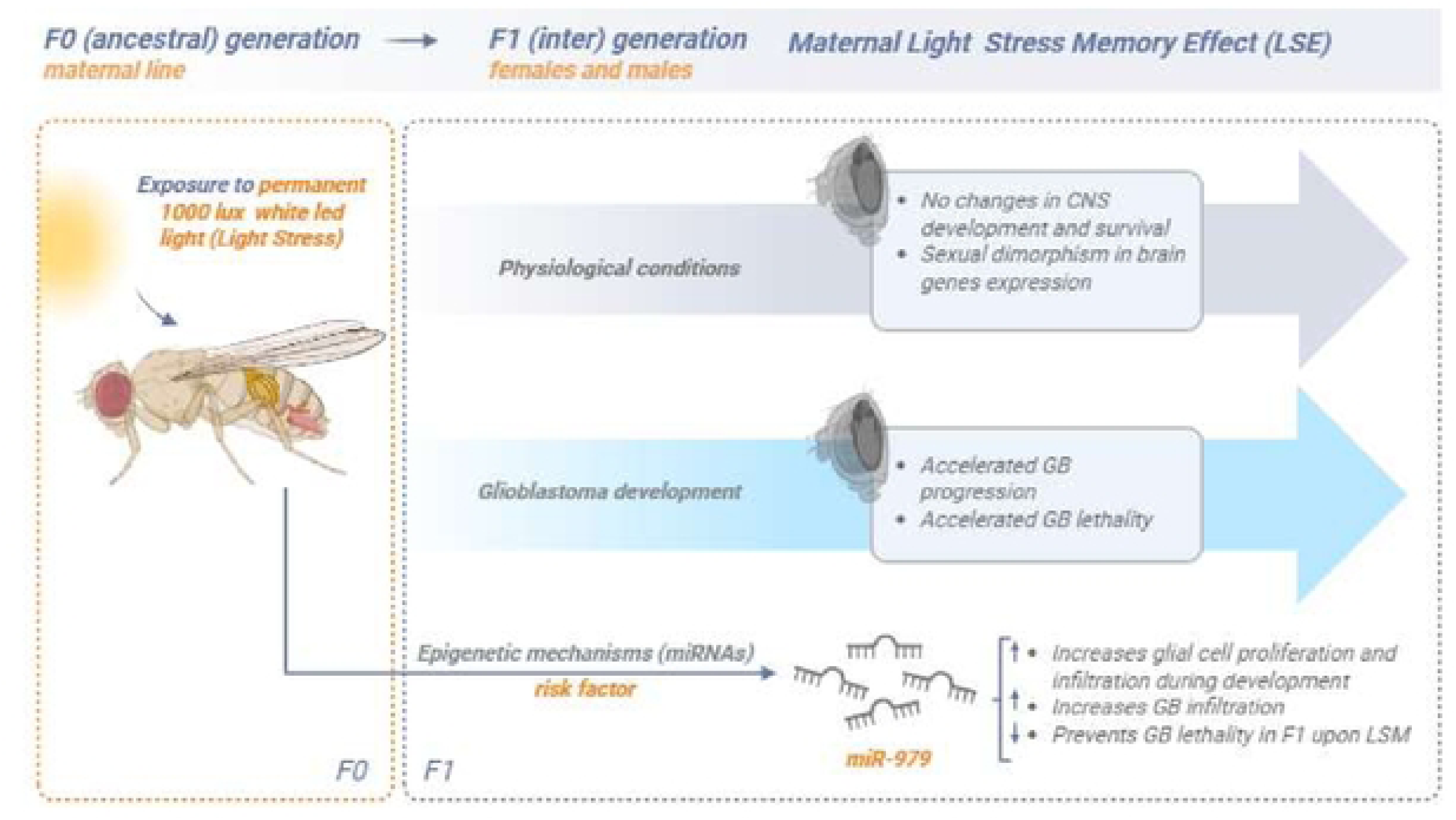

## Introduction

Glioblastoma (GB) is the most common and lethal glioma, classified as a grade IV diffuse astrocytoma. Highly proliferative, invasive and resistant to treatment, it infiltrates surrounding the brain parenchyma^1^ Its incidence is about 3 per 100,000/year^2,3^ and its 5-year survival rate is 6,8%^4^. Due to its diffuse nature, complete resection is impossible but standard therapy includes Temozolomide (TMZ) and radiotherapy to prevent tumor ^5^, but relapse is inevitable, often with a more aggressive phenotype and poor prognosis^5^. However, the reasons behind the wide variability in life expectancy among GB patients—ranging from weeks to years^6^—even among those with similar mutations, remain poorly understood.

*Drosophila melanogaster* has emerged as a powerful model system to reproduce GB pathology, based on the most common genetic mutations in GB patients as glial-specific co-expression of constitutively active forms of dEGFR and dp110 (EGFR and PI3K orthologs respectively in humans), using repo-Gal4 glial-specific transcriptional driver^7^. This *Drosophila* model recapitulates the genetic, cellular, and physiological characteristics of GB in patients, as it reproduces glial proliferation and invasion and the alteration of signaling pathways involved in glioma-neuron communication triggering tumor growth ^8–12^. Moreover, environmental stimuli as light can restore GB growth and lethality in flies^13,14^. A recent study in *Drosophila* demonstrated that the external light-dark input modulates GB features including neurodegeneration and delays tumor progression even after tumor onset through circadian rhythm re-adjustment^13^. These results sustained the findings in mice models exposed to chronic jet lag. Khan et al. described a link between the onset of a primary brain tumor with a circadian link disrupted by GB-related genes^15^. Hence, external factors should be considered in GB study.

Potential lifestyle factors contributing to glioma burden have been explored^16^. Among them, chronic exposure to light, or light pollution has emerged as a relevant environmental stressor, linked to behavioral disruptions, neurodegenerative diseases, increased cancer risk, and tumor aggressiveness^13,17^. Building on Caspi and Moffitt’s 1999^18^ theory of genetic moderation, we propose the *’Brain fitness’* hypothesis: different brains exposed to similar insults show different fitness due to ancestral lifestyle or stress exposure, sensitizing the progeny to pathological outcomes^19,20^.

Environmental stressors contribute to disease through lifelong exposure, from prenatal stages to adulthood, including lifestyle habits (diet, smoking, alcohol, physical activity), occupational exposure, and broader environmental factors like UV radiation, air pollution, or heavy metals^21^. This modern conceptualization of stress-disease contribution is known as *exposome* or *pollutome* ^22,23^. The Developmental Origin of Health and Disease (DOHaD) theory links early-life environmental influences at the time of fertilization, embryonic, fetal, or neonatal stages with genetic intrinsic factors^24^ triggering epigenetic modifications. However, this *fetal basis of adult disease* theory should be expanded to pre-gestational stages when environmental factors can alter the epigenetic status of germ line cells, compromising the fitness of next generations ^25–29^. For instance, work occupation including benzene-related occupational exposures (exposure to paints, solvents or exhaust fumes) or magnetic fields have been linked with an increased CNS cancer or leukemia predisposition in the next generation^30–32^. Therefore, brain cancer should also be considered within the context of environmental and lifestyle factors.

Besides neurological pathologies, brain cancers are increasingly linked to environmental stressors^25,33–36^, including artificial light exposure. Modern lifestyles, have raised exposure to artificial light (light pollution) due to the use of electronic devices based on light-emitting diodes (LEDs), which is a stress source for animals^37^, disrupts circadian rhythms and increases cancer risk^13,15^. Light pollution is associated with shorter lifespan, accelerated aging, locomotor impairment, neurodegeneration, oxidative stress^38–40^, and enhanced tumor traits such as cell growth and migration^41–44^. In *Drosophila melanogaster*, chronic white light exposure shortens lifespan, accelerates aging and neurodegeneration, even in genetically blind flies^39,40,45^. However, blue light particularly affects to metabolic reprogramming, reduces survival, promotes differentially expressed genes in the heads rather than the whole fly body and suggests that the eye/brain are the most affected tissues by blue light^46^. However, the generational effects of parental light exposure remain largely unknown.

Epigenetic generational inheritance refers to epigenetic changes in the germline that are transmitted to subsequent generations in the absence of a permanent environmental exposure across species (McClintock, 1953)^47–58^. These changes involve mechanisms like DNA methylation, histone modifications, and small non-coding RNAs (e.g., microRNAs and siRNAs), which lead to gene expression changes shaping offspring phenotype across different species^59–70^. Small non-coding RNAs (microRNAs and siRNAs) are proposed as maternal transgenerational inheritance vectors across generations in several species of nematodes (*C.elegans*), insects (*D. melanogaster*) and mammals^71–75^. Environmental conditions affect the biogenesis of miRNAs and can be transferred through germ line cells to the next generation^73–76^. For example, in *Drosophila*, a parental Western diet causes transgenerational changes on brain proteome and microRNAs modulation, affecting offspring feeding behavior^77^. In mammals, parental stress alters offspring traits via germline transmission^78–81^. Notably, miRNAs are emerging as epigenetic biomarkers for disease risk, diagnosis, and prognosis, including in neurodegeneration, psychiatric disorders, and cancer^20,82^.

We introduce the concept of *Brain Fitness* as a novel framework to describe the intrinsic capacity of brain cells to respond effectively to injury, pathology, or disease. This state of resilience and adaptability is determined by both intrinsic factors, such as genetic programs and cellular homeostasis, and extrinsic factors, including environmental stimuli and systemic signals. At the molecular level, *Brain Fitness* is defined by the expression of specific gene networks that regulate neuroprotection, plasticity, and repair mechanisms. Importantly, this state is dynamic and can be modulated by epigenetic factors, enabling external influences to shape gene expression patterns and, in certain cases, allowing the transmission of adaptive or maladaptive traits across generations. Research across species indicates that this epigenetic memory is stress-specific, suggesting a potential biomarker of ancestral lifestyle, linking past exposures to inherited disease susceptibility^67,83^. This novel concept provides a dynamic perspective on the ability of brain cells to maintain functional integrity under stress and highlights potential avenues for therapeutic intervention.

Lifestyle factors—diet, behavior, physical activity, occupation, substance use, and stress—have been shown to induce epigenetic changes that persist through development and can be passed to offspring^78–80,84–87^. This evolutionary conserved mechanism supports a suitable adaptation response and a specific genetic program in a changing environment. Hence, it is suggested that modern human exposures might be a cause or contributing factor in the onset of disorders altering individual fitness^88,89^.

The term *memory* in the field of *transgenerational transmission* of stress has been used in plant literature to refer to the process through which information regarding a past stress cue is retained in an organism exposed to this stress for more efficient response to future exposure^90^. However, to avoid potential controversy or ambiguity associated with this term, in the present work we use the expression stress effects instead of stress memory to refer specifically to the consequences of parental exposure to a stressor as observed in the F1 generation when it encounters the same stimulus.

Exposure of parental lines to stressful conditions can alter the vulnerability of the offspring to develop multiple pathological conditions^87^. This corresponds to a Lamarckian evolution perspective wherein an organism can transfer environmental characteristics to the offspring as an accommodation or phenotypic plasticity to past environments, determining gene expression and altering its fitness. Therefore, the DOHaD theory should be amplified to pre-conceptional stages along with the ancestral lifestyle or environmental influences to which they have been exposed^87,91,92^ and we propose the *Brain fitness* hypothesis. Here, we examine the effects of maternal luminic stress on the molecular fitness of their offspring, the generational lasting effects, and the imprinted vulnerability to worse prognosis of GB.

## Results

### Light exposure alters germ line oxidative stress and it impacts on the next generation

Light is one of the key players synchronizing organisms based on day/night periodicity. In the case of *Drosophila*, disruptions in this cycle can alter developmental features, sleeping time, aging, or oxidative levels^40,93^. To investigate whether the luminic stress (LL) protocol optimized in the laboratory has an impact on the progeny, we first exposed 3-4 days-old virgin adult females to a 7-day LL 24 protocol (1,000 luxes white light 24h during 7 days) under standard diet and temperature (25°C) to analyze the germ line oxidative stress (Fig.1A). The control flies were kept in LD 12:12 conditions (12h of light/12h of darkness each day) under standard diet and temperature (25°) (Fig.1A) (Table 1).

**Table 1.** Experimental designed conditions to evaluate maternal light stress protocol and generational effect.

| Maternal light stress experimental settings |
| --- |
| **F0 females** |
| F0 females LD 12:12 (L/D) |
| F0 females LL 24 (L/L) |
| **F1 flies** |
| F1 control LD 12:12 F0 females LD 12:12 (L/D) |
| F1 control LD 12:12 F0 females LL 24 (L/L) |

LL- light stress; LD-light darkness

**Figure 1.**
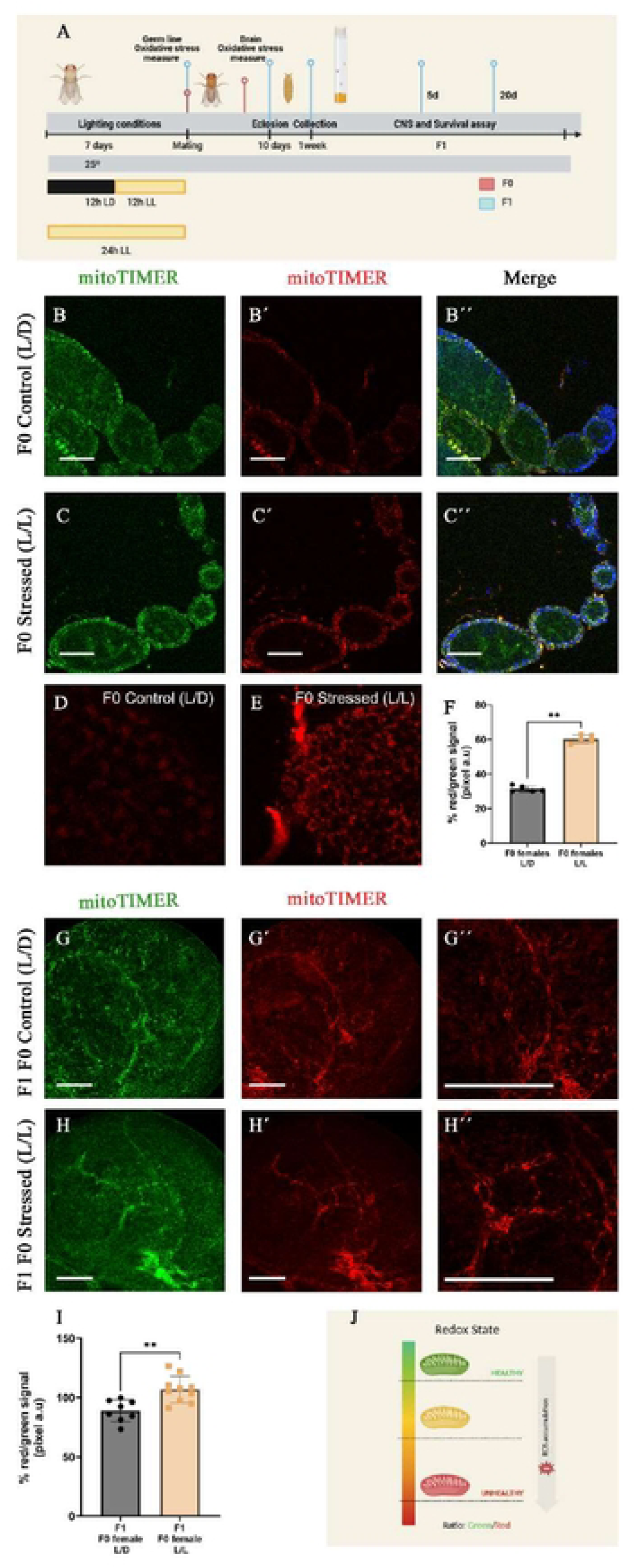
Maternal luminic stress exposure induces changes in F1 CNS oxidative stress through germ line oxidation status. A. Protocol diagram. 0-3 days old virgin females were exposed to different lighting conditions (12:12 L/D, 12:12 L/L) for 7 days and then mated with males to obtain de F1 generation non-exposed to light stress (12:12 L/D). B-Cúú63x confocal mi croscopy i mages of F0 D rosophi I a adult vi rgi n femal e ovari es carry i ng M itoT i mer reporter tool (tub-Gal4-LL7> UAS-Mitotimer) under (B-Búú) luminic control conditions (L/D) and (C-Cú) permanent light exposure or stressed F0 flies (L/L). Fluorescent green signal reports a healthy mitochondria status and turns to red when oxidize in presence of reactive oxygen species. Size bar 100 um. D-E detailed images of red signal of ovaries under both luminic conditions at 63x z3. F. Quantification of pixel intensity signal ratio between red and green signal (a.u arbitrary units) comparing F0 females under luminic control conditions (L/D) and F0 females under a permanent light stimulus (L/L). T-test Student and Mann-Whitnney post-test analyses. ** p<0,005. G-Húú63x confocal microscopy images of F1 Drosophila larva lobe brain from (G-Gúú) F0 females under luminic control conditions (L/D) and (H-H úú) continuous light exposure (L/L), carrying MitoTimerreporter tool (tub-Gal4-LL7>Mitotimer. Gúú Húúdetailed images at 63x z3. Size bar 100 um. I. Quantification of pixel intensity signal ratio between red and green signal (a.u arbitrary units) comparing F1 from females under luminic control conditions (L/D) and F0 females under a permanent light stimulus (L/L). T-test Student *** p<0,001. J. Mitotimer Reporter Tool diagram.

To monitor oxidative stress, we used the reporter MitoTimer that encodes a mitochondria-targeted green fluorescent protein sensitive to oxidation as a reporter for healthy mitochondria^94^, under the control of a ubiquitous actin promoter (*act-gal4 LL7*). Upon oxidative stress, the signal turns red, reporting cumulative redox mitochondrial history^94^(Fig.1J). We analyzed the ovaries of female flies exposed to LL 24h compared to LD 12:12 flies (Fig. 1B-E) (Table 1) and our results described an increase of fluorescent signal towards red upon luminic exposure (Fig. F). Thus, a disruption of circadian LD 12:12 period such as exposing flies to a 24h permanent light period induces mitochondrial oxidative stress in cells of the reproductive system.

Using the same experimental parental LL protocol, we then evaluated the impact on the next generation. In this case, virgin females with the ubiquitous *actgal4 LL7; Tub Gal80 ^TS^* driver were exposed to LL 24h and mated with males carrying the *MitoTimer* tool. F1 flies were kept at LD 12:12 lighting period and 29°C during embryonic development to activate the *Gal80 ^TS^/Gal4-UAS* expression system and F1 3^rd^ instar larvae brains were dissected to measure MitoTimer fluorescence signal (Fig. 1G-H). We quantified the green and red fluorescence signal in F1 brain samples from control-F0 unstressed females and LL-F0 stressed females (Fig. 1 I) (Table 1). Our results showed an increase in mitochondrial oxidative stress in the brains of progeny of LL-F0 stressed females (Fig. 1 I) (Table 1). Thus, the effect of maternal environmental light stress is inherited to the offspring through the maternal germ line leading to an imbalance in the F1 brains’ oxidative stress. Hence, we refer to this phenomenon of luminic environmental stress which triggers a hallmark in the offspring as <u>L</u>ight <u>S</u>tress <u>E</u>ffects (LSE).

### LSE does not alter neural development

We next investigated if the maternal exposure to LL affects the development of the nervous system in the offspring. First, we quantified the number of active zones (i.e., synapses) at the neuromuscular junction (NMJ) in F1 females and males from LD-F0 unstressed females and LL-F0 stressed females of young adult flies (5 days) and old adult flies (20 days) (Fig. 2) (Table 2). Moreover, we quantified the volume of glial membrane in F1 brains. To evaluate the impact of LSE shaping *Brain fitness*, we choose to study young (5 days) and old (20 days) adult flies (Fig. 2).

**Table 2.** Experimental designed conditions to evaluate LSE during neural development.

| Maternal light stress experimental settings |  |
| --- | --- |
| F1 males | F1 females |
| F1 control LD 12:12 F0 females LD 12:12 (L/D) | F1 control LD 12:12 F0 females LD 12:12 (L/D) |
| F1 control LD 12:12 F0 females LL 24 (L/L) | F1 control LD 12:12 F0 females LL 24 (L/L) |

**Figure 2.**
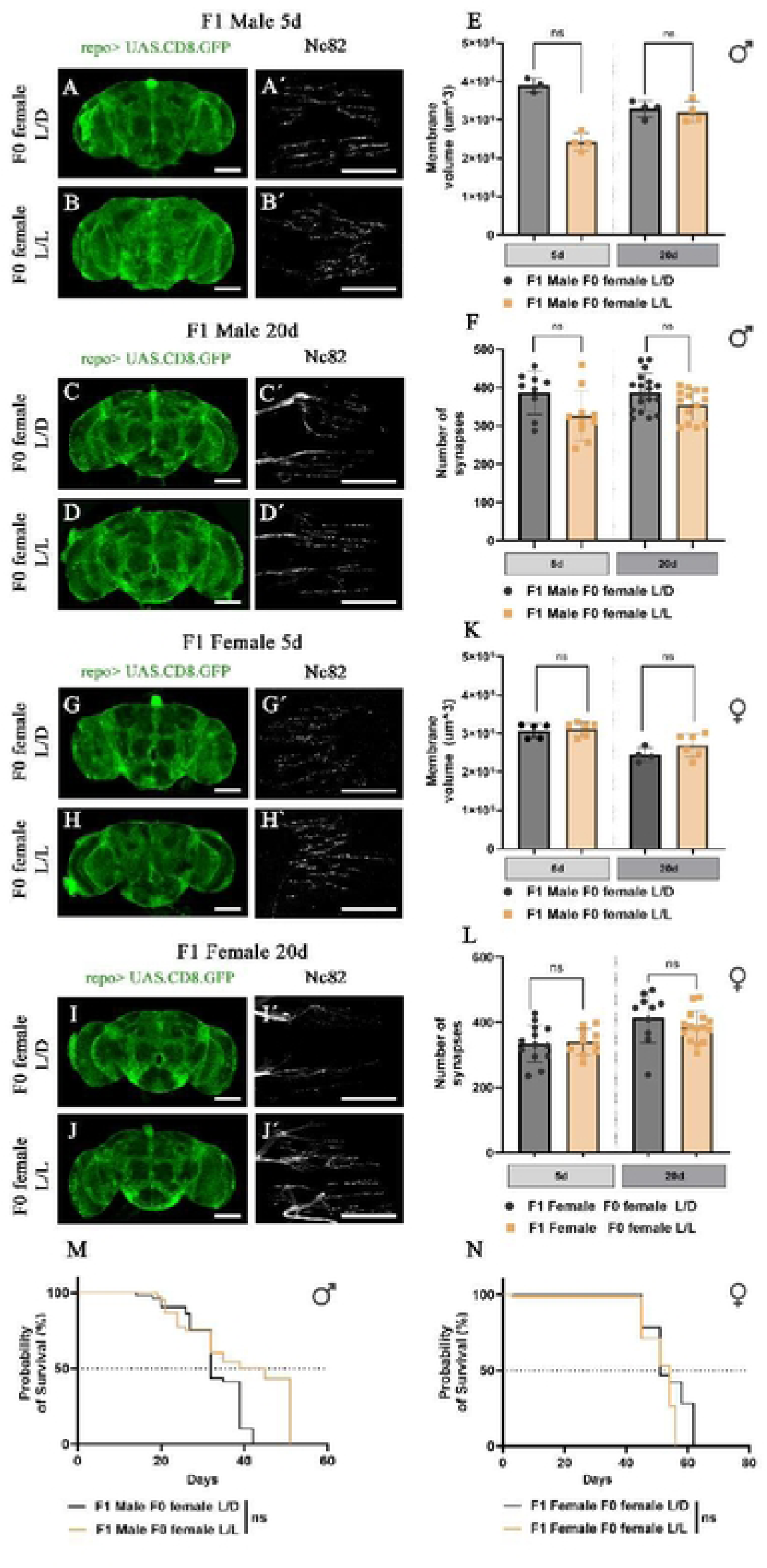
LSE does not impact in F1 offsprings C NS development and survival. A-B, C-D. Confocal microscopy images of F1 Drosophila male adult brains carrying repo> UAS.CD8.GFP construct from (A, C) F0 females under luminic control conditions (L/D) and (B,D) permanent light exposure (L/L) at 5 and 20 days after eclosion. Glial membrane is marked in green color. Size bar 100 urn. Aú·Bú CúDú 20x confocal microscopy images of F1 Drosophila male adult N MJ. A ctive zones are stai ned wi th B ruchpi I ot (Nc82) anti body. E. Quantificati on of membrane volume and F. synapse number comparing F1 males from F0 females under luminic control conditions (L/D) and F0 females under continuous light stimulus (L/L) at 5 and 20 days after eclosion. T-test Student and Mann-Whitnney post-test analyses. ** p<0,005. G-H, I-J. 20x confocal microscopy images of F1 Drosophila female adult brains carrying repo> UAS.CD8.GFP construct from (G, I) F0 females under luminic control conditions (L/D) and (H-J) pemanent light exposure (L/L) at 5 and 20 days after eclosion. K. Quantification of membrane volume and L. synapse number comparing F1 males from F0 females under luminic control conditions (L/D) and F0 females under continuous light stimulus (L/L) at 5 and 20 days after eclosion. T-test Student and T-test Student with Mann-Whitnney post-test analyses. ** p<0,005. M. Graph showing a survival assay of F1 males repo> tubGal80TS; UAS.nryrRFP CS from unstressed F0 females(F1 males control F0L/D), from stressed F0 females (F1 malescontrol F0 females L/L). N. Graph showing a survival assay of F1 females repo> tubGal80TS; UAS.rryrRFP CS from unstressed F0 females (F1 females control F0 L/D), from stressed F0 females (F1 females control F0 females L/L). Statistical analysis included (Mantel-Cox test).

The results show that F1 young adult male flies from LL-F0 stressed females at 5 days after eclosion suffered a decrease in the volume of the glial membrane, while there were no differences in aged flies 20 days post-eclosion (Fig. 2 A-B,E). In addition, F1 males from LL-F0 stressed females did not show a significant decrease in the number of synapses in any selected developmental time (Fig. 2C-D,F). Similarly, in F1 females from LL-F0 stressed females, glial membrane volume or synapse number did not show any difference neither at 5 days nor 20 days of age (Fig 2G-L). Thus, we concluded that LSE does not cause significant alterations in CNS development in the progeny despite the observed brain oxidative stress observed in early development (Fig. 1).

### LSE is not sufficient to elicit reduced survival in non-tumoral conditions, but it accelerates GB progression

We next asked if the LSE is sufficient to sensitize F1 brains for future neurodegenerative processes as those elicited by GB growth. We first performed a survival assay under non-tumoral conditions (Fig. 2M-N) and inducing a CNS aggression such as GB development (Fig. 3A, 3C) (Table 3). We designed the survival experiments with the following non-tumoral conditions: 1) F1 flies under non-tumoral conditions from LL-F0 stressed females and from 2) LD-F0 unstressed females (Fig. 2 M-N); and with the following tumoral conditions: 1) F1 GB flies from LL-F0 stressed females, 2) F1 GB flies from LD-F0 unstressed females and control (non-GB) flies from LD-F0 females (Table 3) (Fig. 3).

**Table 3.** Experimental designed conditions of non-tumoral and tumoral flies to evaluate LSE under pathological conditions.

| Non-tumoral conditions settings |  |
| --- | --- |
| F1 males | F1 females |
| F1 control LD 12:12 F0 females LD 12:12 (L/D) | F1 control LD 12:12 F0 females LD 12:12 (L/D) |
| F1 control LD 12:12 F0 females LL 24 (L/L) | F1 control LD 12:12 F0 females LL 24 (L/L) |

| Tumoral conditions settings |  |
| --- | --- |
| F1 males | F1 females |
| F1 GB LD 12:12 F0 females LD 12:12 (L/D) | F1 GB LD 12:12 F0 females LD 12:12 (L/D) |
| F1 GB LD 12:12 F0 females LL 24 (L/L) | F1 GB LD 12:12 F0 females LL 24 (L/L) |
| F1 control LD 12:12 F0 females LD 12:12 (L/D) | F1 control LD 12:12 F0 females LD 12:12 (L/D) |
| F1 control LD 12:12 F0 females LL 24 (L/L) | F1 control LD 12:12 F0 females LL 24 (L/L) |

**Figure 3.**
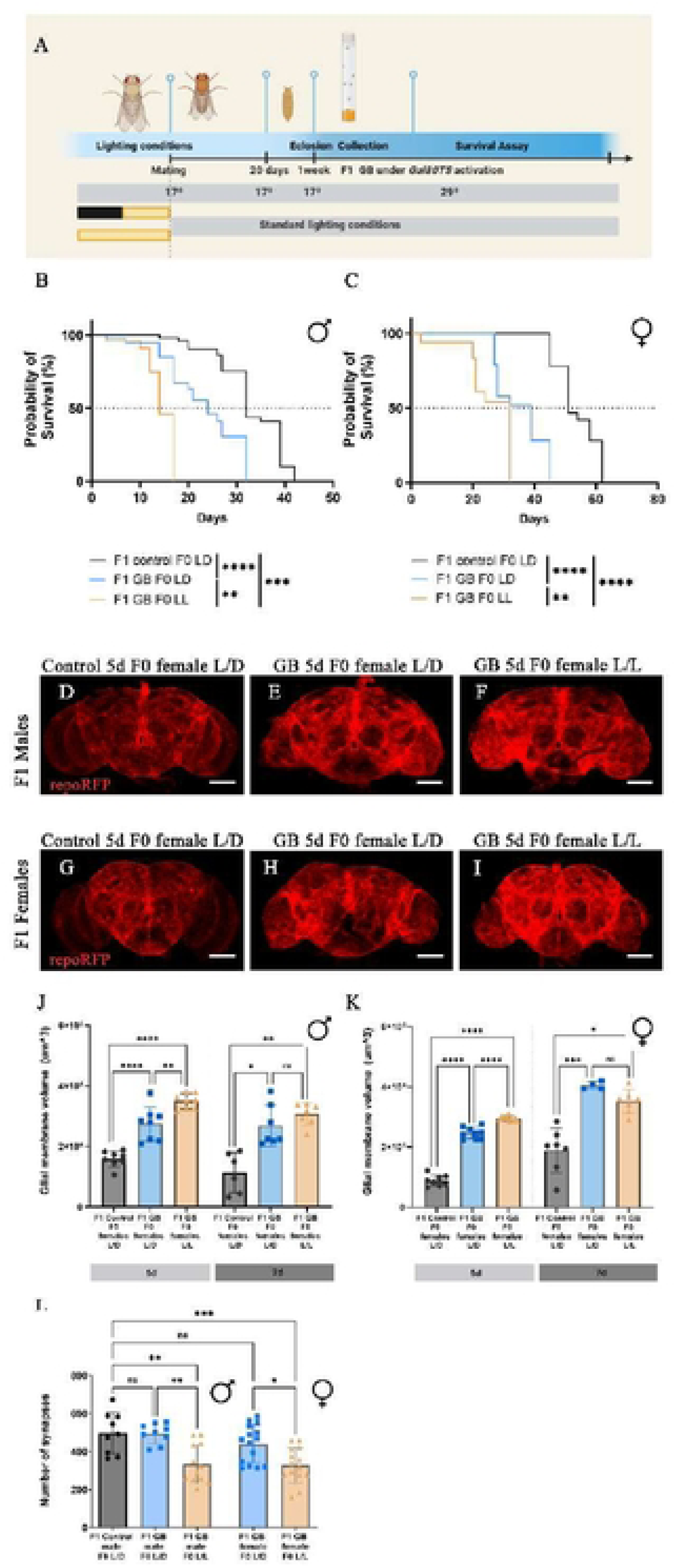
LSE accelerates tumor progression and lethality in F1 progeny. A. Protocol diagram.0-3 days old virgin females were exposed to different lighting conditions (12:12 L/D, 12:12 L/L) for 7 days and then mated with males to obtain de F1 generation non-exposed to light stress (12:12 L/D). F1 progeny were developed at 17ł until eclosion that they were transferred to 29ł to induce a tumor under the control of repo> tubGal80TS, UAS.-rryrRFP genetic construct. B-C. Graphs showing survival assay of F1 males (B) and females (C) developing a tumor under different maternal lighting conditions: F1 control F0 LD (repo-Gal4> tubGal80TS; UAS-rryrRFP construct from unstressed F0 females, F1 GB F0 LD (repo-Gal4> tubGal80TS, UAS-dEGFRZ, UAS-dp110CAAX from F0 females under luminic control conditions (L/D)) and F1 GB F0 LL (repo-Gal4> tubGal80TS, UAS-dEGFRZ, UAS-dp110CAAX from continuous light exposure (L/L)). Statistical analysis included (Mantel-Cox test) ** p<0,005, *** p<0,001, **** p<0,0001. D-I. Confocal microscopy images of F1 Drosophila males (D-F) and females(G­I) adult brains at 5 days developing a tumour. Glial membrane is marked in red. Size bar 100 um J-K. Quantification of membrane volume at 5 days and 7 days of tumor development in F1 males (J) and F1 females (K) One-way ANNOVA tes, Bonferroniii multiple comparition pos-tes, and One-way Kruskal-Wallis test Dunnii multiple comparition post-test * p<0,05„ ** p<0,005, *** p<0,001. L. Quantification of synapse number. One way ANNOVA, Bonferroniíε multiple comparition post-test is used * p<0,05.,** p<0.05, *** p<0,001.

The results show that control F1 males from stressed females did not change their survival rates respective to control males from LD-F0 unstressed females (Fig. 2M). Similarly, F1 females from LL-F0 stressed females did not show reduced survival under LSE (Fig. 2N). Thus, although we observed a sensitized brain in the progeny of the females that were exposed to an environmental luminic aggression (LL 24h) in terms of oxidative stress, this pre-conditioned brain does not alter neural development nor individual survival under non-tumoral conditions.

Next, we analyzed the survival of flies bearing GB development under maternal light stress conditions (LD 12:12 vs LL 24h) (Fig. 3). Glioma flies were kept at 17° during development until pupal emersion and adult gathering to initiate the experimental protocol at 29° to activate genetic machinery to induce the tumor (Fig. 3A). We observed that F1 flies developing a GB facing LSE showed premature death compared to F1 GB flies under control maternal lighting conditions (Fig. 3B-C). The described effects were reproduced in both male (Fig. 3B) and female (Fig. 3C) F1 flies. Survival differences were also described comparing both F1 GB conditions with F1 control flies (Fig. 3B-C). These results suggest that LSE effect on the progeny could be initially hidden in the absence of malignancy development, but it reveals on the individual fitness under a pathological progression. In the case of GB, the progeny of stressed females shows premature death, therefore, LSE seems to sensitize the F1 brain to this disease.

### LSE accelerates tumor invasion in the progeny

To elucidate the mechanisms through which LSE could worsen the GB phenotype in the progeny, we quantified GB infiltration capacity through the expansion of tumor microtubes (TMs) by quantifying glial membrane volume^13,95–97^ after 5day tumor development (Fig. 3D-I). Thus, we studied tumor invasion in F1 females and males from LL-F0 stressed females, LD-F0 unstressed females and F1control flies from LD-F0 unstressed females (Table 4).

**Table 4.** Experimental design conditions of tumoral flies to evaluate LSE effect on GB features.

| Experimental settings |  |
| --- | --- |
| F1 males | F1 females |
| F1 GB LD 12:12 F0 females LD 12:12 (L/D) | F1 GB LD 12:12 F0 females LD 12:12 (L/D) |
| F1 GB LD 12:12 F0 females LL 24 (L/L) | F1 GB LD 12:12 F0 females LL 24 (L/L) |
| F1 control LD 12:12 F0 females LD 12:12 (L/D) | F1 control LD 12:12 F0 females LD 12:12 (L/D) |

LL- light stress; LD-light darkness; GB-Glioblastoma

We observed an increase of glial membrane volume upon GB induction in the progeny of any condition developing a tumor (Fig. 3D-I) (Table 4). F1 GB males (Fig. 3D-F) and females (Fig. 3G-I) from LL-F0 stressed females showed an additional increase in the glial membrane volume compared with F1 GB flies from LD-F0 unstressed females. As expected, LSE in F1 flies leads to an increase in the tumor invasion ability upon 5 days of tumor development (Fig. 3J-K). Furthermore, we quantified tumor volume at 7 days to trace tumor development at advanced tumoral stages (Fig. 3J-K). Our results showed that tumoral glial volume differences are not maintained in GB samples at 7 days of tumor development (Fig. 3J-K). Therefore, LSE influences the brain’s susceptibility to tumor growth and aggressiveness from the early stages of tumor development (Tables 3-4). As a result, GB progression is accelerated under LSE conditions.

### LSE accelerates tumor proliferation in the progeny

To determine the effect of LSE in tumor progression, we quantified the number of glial cells ^13,95–97^ in GB and control flies under different environmental maternal light stress conditions as described in Table 4. Consistent with tumor invasion, the number of glial cells in F1 males and females bearing a GB from LL-F0 stressed females is significantly higher compared with F1 GB flies from LD-F0 unstressed females (Supplementary 1 A-B).

The results show that 5 days of tumor development is not enough to provoke significant increase in GB cells number in F1 GB flies from LD-F0 unstressed females compared with F1 control flies from LD-F0 (Supplementary 1 A-B). However, in line with the invasion results, GB brains from LL-F0 stressed females showed an increase in GB cell number (Supplementary 1 A-B). In addition, we observed that after 7 days of tumor development the quantifications showed glial cell number differences (Supplementary 1 A-B). Based on these results, we propose that LSE pre-conditioned the brain in the next generation, increasing its sensitivity such that, in the context of a disease like GB, pathological progression is accelerated.

### LSE accelerates GB-induced neurodegeneration

Since synapse number reduction is one of the first steps in a neurodegenerative process including GB ^95–97^, we quantified synapse loss associated with tumor growth at various GB developmental stages to determine whether LSE influences this process (Fig. 3J). Notably, at 5 days of GB development, F1 GB males and females from LL-F0 stressed females or LD-F0 unstressed females did not show a reduction in the number of synapses compared to F1 control flies from LD-F0 unstressed females (Fig. 3J). These results suggested that 5 days of tumor development are not enough to promote tumor-induced synapse number reduction.

Next, we quantified synapse numbers at 7 days of GB development. We observed that F1 males and females from LL-F0 stressed females that develop a GB showed a reduced number of active zones compared with F1 GB flies from LD-F0 unstressed females (Fig. 3M). Although 7 days of tumor development are not enough to promote GB associated synapse number reduction, LSE sensitizes the brain, and it accelerates synapse loss. We expected that LSE will accelerate TMs formation as the first step in tumor growth, and glial proliferation at early stages (5 days), while it not being sufficient to trigger neurodegeneration. However, we found that, at 7 days of tumor development, TMs expansion differences due to LSE are not significant while GB cell number and synapse number reduction are evident. Taken all together, LSE progressively alters tumor growth and progression in F1 flies leading to accelerated key, but not all, malignancy features (Table 5).

**Table 5.**
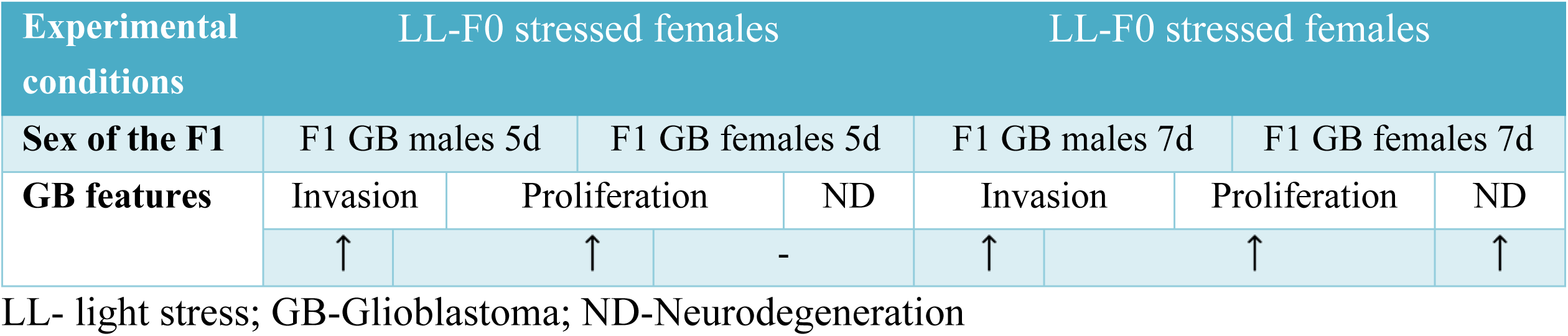
GB features quantification in F1 brains from LL-F0 stressed females at 5 and 7 days of tumor development vs normal-F0 unstressed flies.

### Stress-type and sex-specific genetic profile

Environmental stress exposure induces changes in the transcriptomic profile across different species^98–101^and it also displays a generational effect on the offspring’s genetic program with a sex bias that impacts individual fitness ^102–106^. To address how parental environmental conditions alter their offspring’s brains’ genetic profiles, we studied differences in gene expression in F1 samples coming from F0 females exposed to different environmental stressors. We performed RNAseq analysis for F1 males and females whose mothers had been exposed to luminic or dietary stress (HFD, high fat diet and SD standard diet)^107,108^ compared to control conditions (F1 control flies from LL-F0 stressed females, F1 control flies from LD-F0 unstressed females, F1 control flies from HFD-F0 females, F1 control flies from SD-F0 females).

There are 4681 genes with altered expression in F1 males descending from F0 females who have been exposed to LL compared to F1 males from F0 LD control females. Of these, 3088 (65.97%) are also altered in F1 females from F0 LL females compared to controls. However, these F1 females descendant from F0 LL females have an additional 2087 genes (for a total of 5175 genes) that show different expression as a result of F0 LL and do not appear to be changed in F1 males (Fig. 4A).

**Figure 4.**
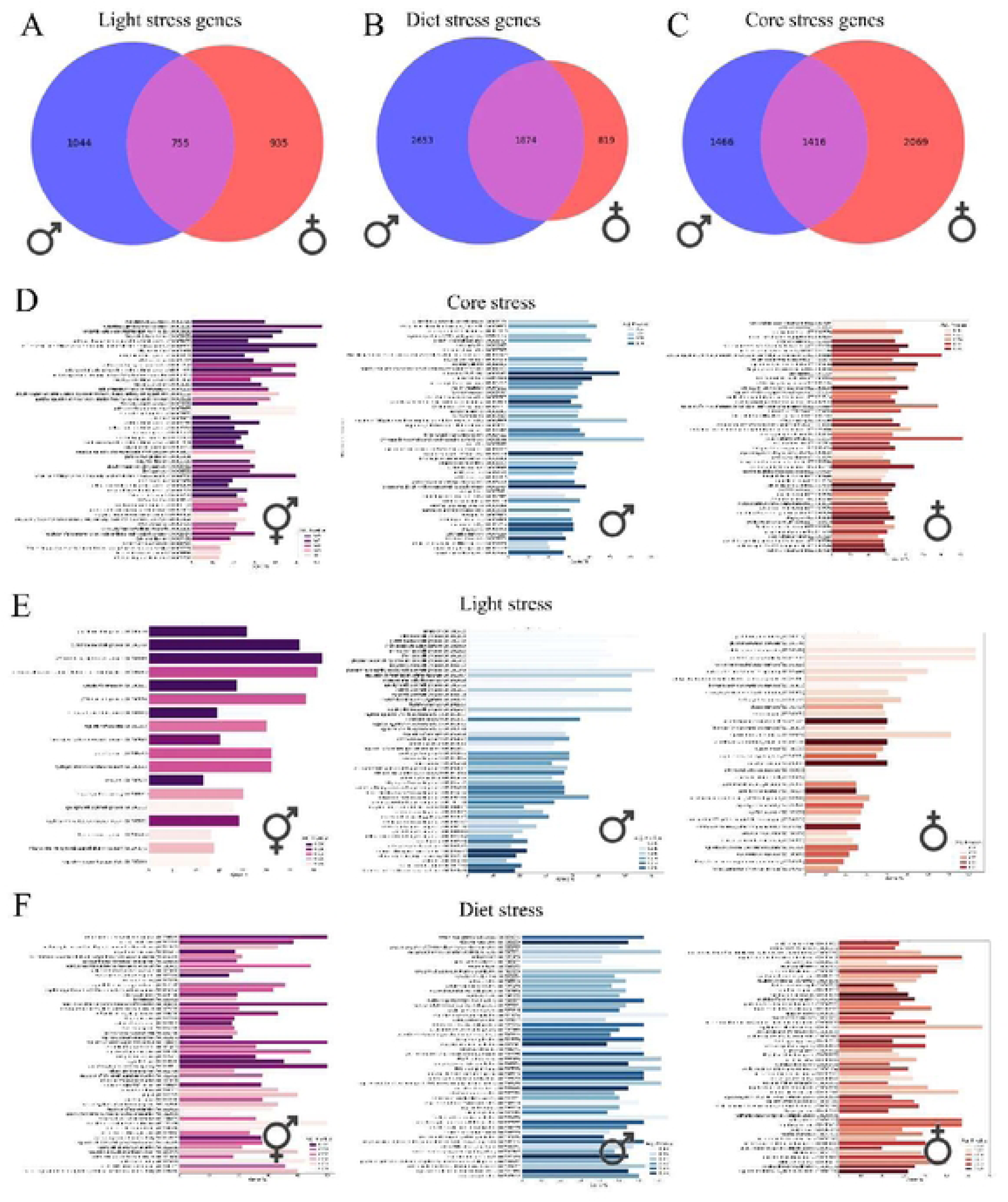
Maternal stress causes alterations in gene expression in an offspring-sex specific and stress-source specific manner. A. Venn diagram of the number of shared, male specific and female specific genes that are altered in F1 flies after maternal exposure to permanent light (Light stress) conditions, compared to controls with maternal L D. B. V enn diagram of the number of shared, male specific and female specific genes that are altered in F1 flies after maternal exposure to high-fat diet conditions (Diet stress), compared to controls with maternal LD. C. Venn diagram comparison of shared, male specific and female specific genes altered in progeny after maternal stress, regardless of the source of the stress. D. Significantly enriched terms in the shared, male-specific and female-specific core stress genelist (p-value< 0.05), found within the library ’GO Biological Processes 2018_. F. Significantly enriched terms in the shared, male-specific and female-specific light stress genelist (p-value<0.05), found within the library ’GO Biological Processes 2018_. F. Significantly enriched terms i n the shared, male-specific and female-specific diet stress genelist (p-value<0.05), found within the library ’GO Biological Processes 2018-.

To probe the specificity of the stress response to the stressor, we performed the same analysis for progeny from high fat diet (HFD) stressed females, as diet can be a source of stress in *Drosophila* and other organisms^79,125,126^. In this case, there are 7409 genes with altered expression in F1 males coming from HFD stressed F0 females compared to controls, compared to 6178 genes altered in F1 females from HFD stressed mothers. In this case, 5230 genes are altered in both F1 males and females, accounting for 84.66% of those altered in F1 females and only 70.59% of those altered in F1 males (Fig. 4B). These results indicate that maternal LS has more of an impact on female progeny than on male progeny, while maternal HFD has the opposite effect.

Accounting for this female bias of the LSE, we investigated mechanisms by which it could be affecting the aggressiveness of tumor growth. We analyzed the contribution of genes associated to GB progression and invasion at early stages of tumor development that precede cellular effects^96,97^. Matrix Metalloproteases (MMPs) participate in tumor progression and TMs network formation that lead to an invasion through the neighboring healthy brain tissue^97^. Consistent with our hypothesis, *Mmp1* expression levels were increased in the brain of F1 females and males (more than 4 and 2 times respectively) from LL-F0 stressed females. However, *Mmp2* expression levels did not show any notable change under any condition. Moreover, genes associated to JNK signaling (*bsk, Tak1, kay, wnd, slpr)* and required for expansion of tumor microtubes in GB^97^ are upregulated in F1 females. Thus, the results suggest that F1 GB phenotype observed in females and males is caused by different mechanisms.

### Inherited stress genetic hallmark

To define a putative gene expression signature of stress, regardless of the nature of the stressor, we next compared the list of LL altered genes to that of HFD altered genes. Of the 4681 genes altered in F1 males upon F0 LL, 61.57% are also altered in F1 males upon F0 HFD. Nevertheless 1799 genes are specific to F0 LL and 4527 are specific to F0 HFD in male progeny. In contrast, of the 5175 genes altered in F1 females after F0 LL, 67.29% of genes are also affected by F0 HFD. In the female progeny, 1690 genes are specific to F0 LL and 2693 to F0 HFD. (Fig. 4C)

Upon further inspection of the gene lists that are not dependent on the source of the stress (non stress-specific), we found that 1416 genes change in both F1 males and females when F0 females are stressed (non-sex and non-stress specific). This accounts for 50.18% of the genes altered in F1 males and 40.63% of the genes altered in F1 females when not considering the nature of the stressor. We analyzed the enrichment in GO terms associated to Biological Processes, in both F1 males and females we found that maternal stress is reflected in changes in gene expression of genes associated to RNA processing (GO: 0030490, 0009451, 0006364, 0001510, etc.), mitochondrial function (GO: 0140053, 0032543), fatty acid processing (GO:0006635, 0019395, 0009062), protein catabolism (GO:0010498, GO:0043161 and GO:0006511), and the ERAD pathway (GO:0036503, GO:0030433), among others. This defines a non-sex specific biological stress hallmark (Fig. 4D).

In enrichment analysis^109^ of the sex-specific but non-stress specific gene lists the results show that maternal stress in F1 males also changes expression of genes related to metabolism (GO:0009152, 0044267, 0006487) and notably of genes associated to neuron development, learning and memory (GO: 0042551, 0007616, 0008306, 0007411, 0007274) as well as RNA processing (GO: 0034470, 0006397, 0000398). For their part, F1 females also show altered expression of genes related to metabolism (GO: 0042593, 0016567) and RNA processing (GO: 0006364, 0030490, 0035194) after maternal stress, regardless of the stressor. Of note, in F1 females there is also significant enrichment of altered genes associated to signaling pathways including Wnt, SREBP and Ras (GO: 0090090, 0032933, 0046578) and to perception of sound, mechanical stimuli and pain (GO: 0007605, 0050954, 0019233) (Fig. 4D).

We repeated this analysis focusing on those genes that are altered only after maternal LL conditions in F1 males and females. In this case 755 genes change only after F0 LL but are not specific to the sex of the progeny, corresponding to 41.97% of all genes that are altered in F1 males and 44.67% of all genes altered in F1 females. The shared genes show enrichment for terms associated with metabolic processes (GO: 0006518, 0006749), charged particle transport (GO: 0015985, 0015992, 1902600, 0055085) and notably eye pigment biosynthesis (GO: 0006726) and cuticle development (GO:0040003). In F1 males of LL F0 mothers, specifically altered genes map to GO terms associated to chitin metabolism (GO: 0006030, 0040003, 0006032, 0006031, etc.), fatty acid metabolism (GO: 0019395, 0006635, 0035337, 0009062, etc.) and bacterium defense (GO: 0042742, 0050829). For their part, F1 females of LL F0 mothers show specific alteration in genes that correspond to GO terms linked to particle transport (GO: 0015986, 0015985, 0098662, 0015758, etc.), response to ROS (GO: 0000302, 0006801, 0019430), cell-cell adhesion (GO: 0044331) and muscle assembly and function (GO: 0030239, 0031302, 0006936). (Fig. 4E) When looking at the genes that are specific to maternal HFD stress conditions, we find 1874 genes that are altered in both F1 males and females, constituting 41.40% of the genes that change in F1 males and 69.59% of those that are affected in F1 females. The genes that are common to all the progeny are linked to GO terms that include intracellular transport (GO: 0099003, 0007035, 0046822, 0046825, etc.), signal transduction (GO: 0035023, 0009966, 0046425), neuronal function (GO: 0007129, 0030516) and learning (GO: 0008306) as well as autophagy (GO: 1901096, 0061912). For their part, F1 males of F0 HFD mothers show alteration in expression of genes associated to embryonic development (GO: 0048568, 2000737, 1902875, 0035017), DNA recombination (GO: 0044774, 0006312, 0045910, 0006265, 0016578) and neuron projection (GO: 0010976, 0030516), while F1 females from HFD F0 females mainly show differences in stem cell processes (GO: 0048867, 0007219, 0072091, 0045596) and cell communication (GO: 0010647, 0023056, 0001952). (Fig. 4F)

Taken all together, these data indicate that while there is a common genetic hallmark to biological stress that is independent of the sex of the progeny, there are additional progeny-sex-dependent factors that might also explain the different effects of stress on tumor growth and progression when comparing F1 females and F1 males. Plus, certain effects are specific to the source of the stress, with some differences also being dependent on the sex of the progeny.

### *miR-979-3p* as one of the potential epigenetic mechanisms for LSE

To further investigate potential mechanisms that could be mediating the stress effects in the progeny, we performed an *in-silico* analysis by the targeting miRNA predictor *TargetScanFly*^110^. In this way we identified the *Drosophila* miRNA *miR-979-3p*, one of the 12 miRNAs in the *miR-972C* cluster located in the X chromosome^111^, as it targets multiple genes that were altered after parental stress.

Next, to determine the effect of *miR-979* modulation in CNS development we up-regulated *miR-979* expression (*UAS-miR-979*), or we reduced *miR-979* activity (*miR-979 sponge*) in glial cells (*repo-Gal4*) (Fig. 5, Fig.6). We observed that the up-regulation of *miR-979* in glial cells caused the expansion of glial membrane and glial cell number (Fig. 5 A-F), albeit, it did not alter synapse number (Fig. 5 G-I). Thus, the up-regulation of *miR-979* appears to be a risk factor to sensitize glial cells to grow and infiltrate within the brain, similar to the behavior of GB cells. However, up-regulation does not reduce synapse number, as the induction of GB does. In addition, we evaluated whether *miR-979* modulation alters survival rates (Fig.5 J-K) and the results described no impact on male nor female flies’ survival upon miRNA modulation. Then, we tested if *miR-979* could be a risk factor correlating with GB malignancy (Fig. 5 L-M). We first quantified GB infiltration and glial cell number of F1 females and males upon *miR-979* up-regulation in GB cells at 5 days of tumor onset (Fig. 5 L-M). We observed a significantly larger increase of glial membrane volume upon *miR-979* up-regulation than in GB samples, and a synergistic effect of GB and *miR-979* up-regulation (Fig. 5 L-M). The combination of GB and *miR-979* up-regulation describes an accelerated glioma infiltration compared to GB conditions without altered *miR-979*. In the case of glial cell number, we did not observe that the up-regulation of *miR-979* increased glial cell proliferation but it induced an accelerated glial infiltration and growth (Supplementary 1 C-D).

**Figure 5.**
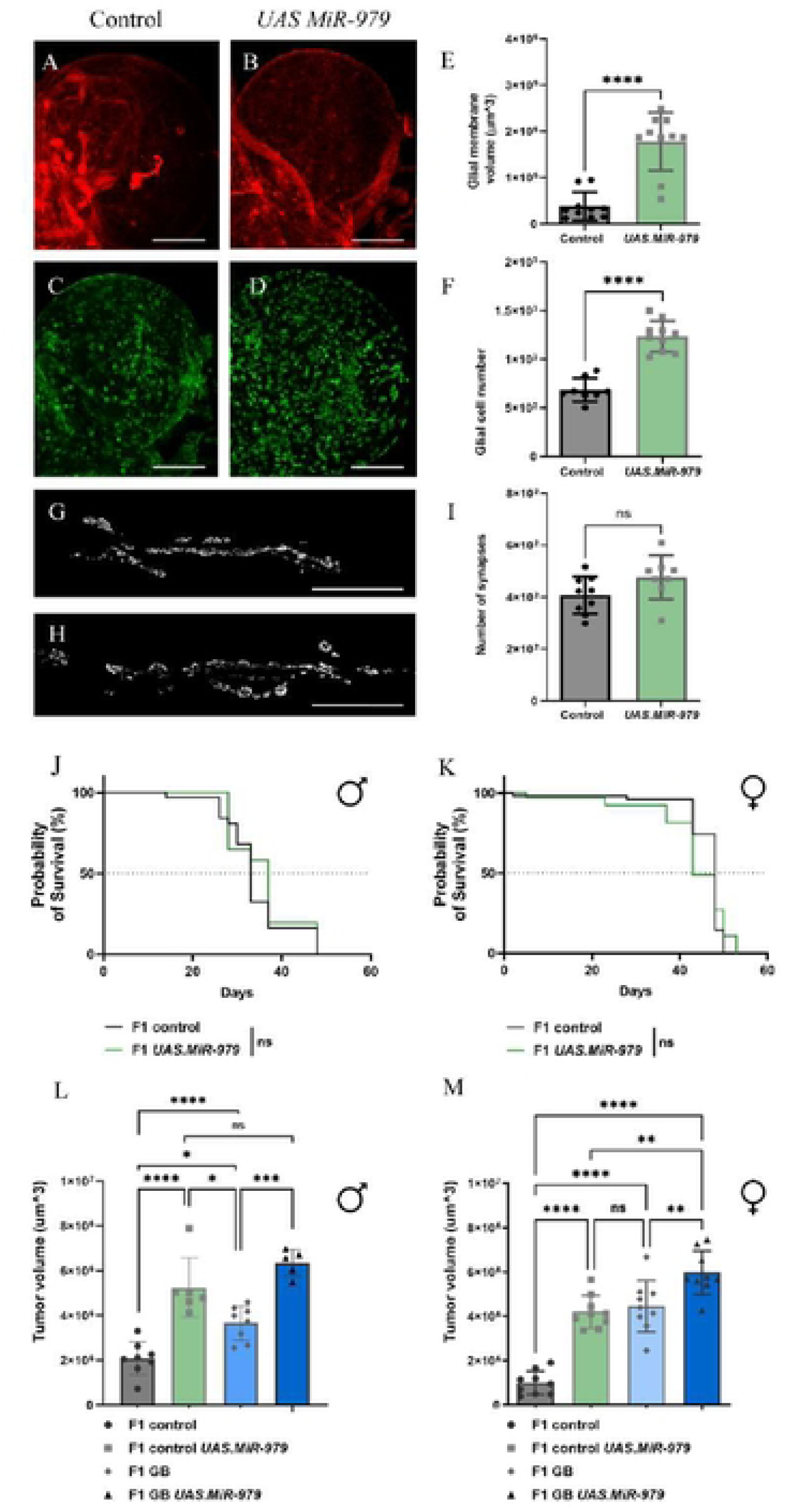
MiR-979 up-regulation alters glial physiology and accelerates tumor progression. A-D. Confocal microscopy images of larval brain lobes and NMJ (G, H) of control flies (repo> LacZ; ihogRFP) and UAS.MiR-979(repo> UAS.MiR-979, ihogRFP). Glial membrane is marked with ihogRFP and glial nuclei are stained with repo antibody in green. Quantification of glial membrane volume(E), number of glial cells (F) and number of synapses (I). T-test Student. **** p<0,0001. Survival graphs of F1 males (J) and F1 females (K) of control flies (repo> LacZ; ihogRFP) and UAS.MiR-979 (repo> UAS.MiR-979, ihogRFP). Statistical analysis included (Mantel-Cox test). Quantification of tumor volume membrane at 5 days of tumor development in F1 males (L) and F1 females (M) of F1 control (repo-Gal4> tubGal80TS CS), F1 control UAS.MiR-979(repo-Gal4> tubGal80TS; UAS.MiR-979), F1 GB (repo-Gal4> tubGal80TS, UAS-dEGFRZ, UAS-dp110CAAX), F1 GB UAS.MiR-979 (repo-Gal4> tubGal80TS, UAS-dEGFRZ, UAS-dp110CAAX; UAS.MiR-979). One-way ANNOVA test, Bonferroniíε multiple compariti on pos-tes, * p<0,05., ** p<0,005, *** p<0,001.

**Figure 6.**
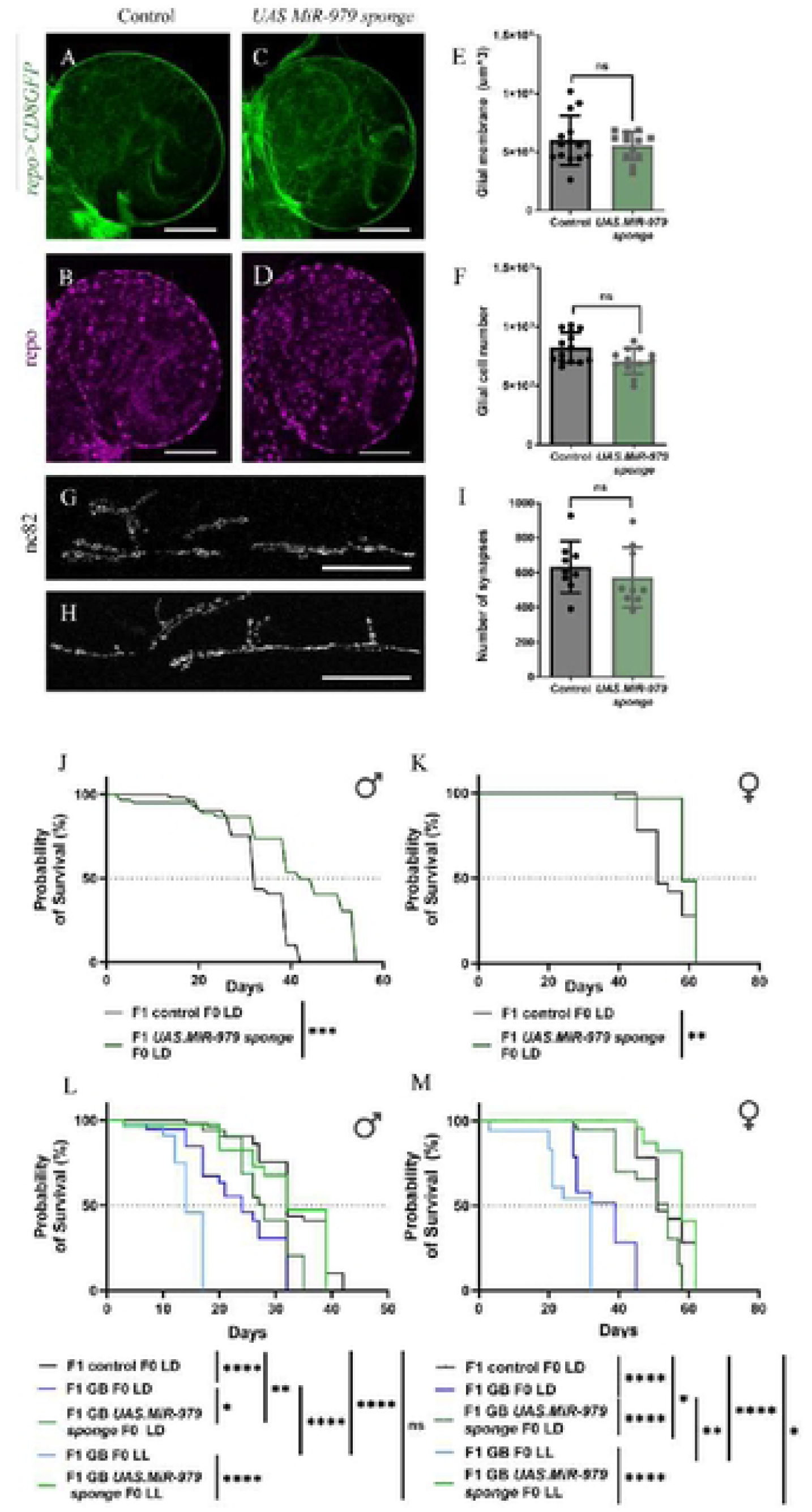
MiR-979 down-regulation prevents tumor lethality. A-D. Confocal microscopy images of larval brain lobes and NMJ (G, H) of control flies (repo> LacZ; UAS.CD8-GFP) and UAS.MiR-979 sponge (repo> UAS.Mi R-979 sponge, UAS.CD8-GFP). Glial membrane is marked with UAS.CD8-GFP, glial cell nuclei are stained with repo anti body in magenta and active zones are stained with Bruchpilot (Nc82) antibody. Quantification of glial membrane volume (E), number of glial cells (F) and number of synapses (I). T-test Student Survival graphs of F1 males (J) and FI females (K) of control flies (repo> LacZ; UAS.CD8-GFP) and UAS.MiR-979 sponge (repo> UAS.MiR-979 sponge, UAS.CD8-GFP). Statistical analysis included (Mantel-Cox test) ** p<0,005, *** p<0,001. Survival graphs of F1 males (L) and F1 females (M) of F1 control LD (F1 repo-Gal4> tubGal80TS CS F0 LD), F1 GB LD (F1 repo-Gal4> tubGal80TS, UAS-dEGFRZ, UAS-dp110CAAX F0 LD), F1 GB UAS.MiR-979 sponge LD (F1 repo-Gal4> tubGal80TS, UAS-dEGFRZ, UAS-dp110CAAX; U AS. Mi R-979 sponge LD), F1 GB LL (F1 repo-Gal4> tubGal80TS, UAS-dEGFRZ, UAS-dp110CAAX F0 LL), F1 GB UAS.MiR-979 sponge LL (F1 repo-Gal4> tubGal80TS, UAS-dEGFRZ, UAS-dp110CAAX; UAS.MiR-979 sponge LL). Statistical analysis included (Mantel-Cox test) * p<0,05** p<0,005, *** p<0,001, **** p<0,0001.

**Figure 7.**
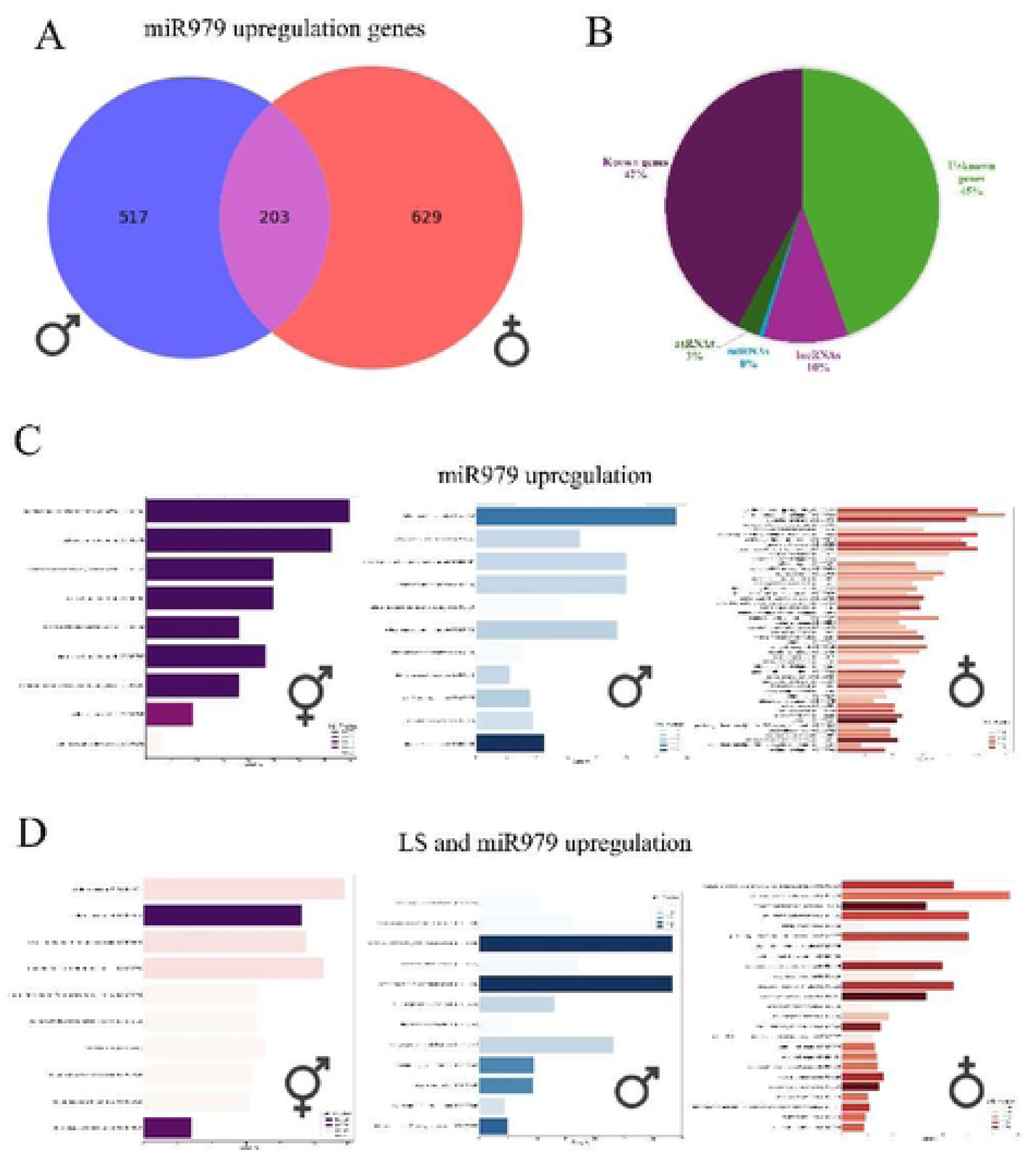
Upregulation of miR979 causes sex-specific alterations that also appear after maternal light stress exposure. A. Venn diagram comparison of shared, male specific and female specific genes altered after upregulation of miR979 B. Average proportion of known genes, undefined genes, IncRNAs, as RNAs and miRN As in the altered genelist C. Significantly enriched terms in the shared, male-specific and female-specific miR-979 upregulation genelist (p-value<0.05), found within the library ’GO Biological Processes 2018_, after filtering out unknown genes D. Significantly enriched terms in the shared, male-specific and female-specific common core of maternal light stress and miR-979 upregulation genelist (p-value<0.05), found within the library ’GO Biological Processes 2018after filtering out unknown genes.

On the contrary, the results show that the loss of function of *miR-979* in glial cells did not alter the glial membrane surface nor glial cell number (Fig. 6A-F). Additionally, *miR-979* sponge in glial cells did not affect synapse number (Fig.6 G-I). Furthermore, we evaluated whether *miR-979* modulation alters survival rates (Fig.6 J-K) and the results show an increased survival rates in male and female flies’ upon miRNA modulation (Fig.6 J-K). Finally, we decided to test the role of *miR-979* combining GB development and maternal lighting conditions (Fig. 6 L-M). We observed that the loss of function of *miR-979* in flies bearing a GB prevents GB early lethality (Fig. 6 L-M). In addition, we also observed this lethality prevention, reaching levels like control flies’, by using the *miR-979* loss of function tool in flies bearing a GB and LSE (Fig. 6 L-M). Hence, we conclude that *miR-979* is a risk factor for GB lethality and its reduced activity inhibits tumor progression that prevent GB flies from an early death.

**Table 6.** Experimental design conditions of tumoral flies to evaluate *miR-979* effect.

| Non-tumoral conditions settings |  |
| --- | --- |
| **F1 males** | **F1 females** |
| F1 control LD 12:12 F0 females LD 12:12 (L/D) | F1 control LD 12:12 F0 females LD 12:12 (L/D) |
| F1 control *UAS-miR-979* LD 12:12 F0 females LD 12:12 (L/D) | F1 control *UAS-miR-979* LD 12:12 F0 females LD 12:12 (L/D) |
| F1 control *miR-979 sponge* LD 12:12 F0 females LD 12:12 (L/D) | F1 control *miR-979 sponge* LD 12:12 F0 females LD 12:12 (L/D) |
| Tumoral conditions settings |  |
| **F1 males** | **F1 females** |
| F1 control LD 12:12 F0 females LD 12:12 (L/D) | F1 control LD 12:12 F0 females LD 12:12 (L/D) |
| F1 GB *UAS-miR-979* LD 12:12 F0 females LD 12:12 (L/D) | F1 GB *UAS-miR-979* LD 12:12 F0 females LD 12:12 (L/D) |
| F1 GB *UAS-miR-979 sponge* LD 12:12 F0 females LD 12:12 (L/D) | F1 GB *UAS-miR-979 sponge* LD 12:12 F0 females LD 12:12 (L/D) |
| F1 GB *UAS-miR-979 sponge* LD 12:12 F0 females LL 24 (L/L) | F1 GB *UAS-miR-979 sponge* LD 12:12 F0 females LL 24 (L/L) |

LL- light stress; LD-light darkness; GB-Glioblastoma

Finally, we performed an RNAseq to elucidate the genes that are modulated by *miR-979* and to establish a link between miRNA levels of expression and the aggressive GB phenotype. In this case we used flies from unstressed F0 females in all conditions and studied the effects of *miR-979* up-regulation by comparing control males and females to flies with expression of the *UAS-miR-979*.

There are 203 genes that are significantly affected by up-regulated *miR-979* in both males and females. These constitute 28 % of the 720 genes that are affected in males and 24 % of the 832 genes that are affected in females (Fig. 6A). This shared list of genes is significantly enriched in GO terms related to polytene chromosome formation (GO: 0035080, 0035079) and protein folding (GO: 0035967, 0006986, 0051084, 0034620, 0051085), confirming the function of *miR-979* as an epigenetic modulator. When looking at the genes that are significantly altered in only males or only females, no GO terms are significantly enriched. These low numbers of enriched GO terms are likely due to a large number of these transcripts being either of undefined function (46 % of non sex specific genes, 44 % of male specific genes and 46 % of female specific genes) or corresponding to lnc:RNAs (7.30% of non sex specific genes, 12 % of male genes and 11 % of female genes), antisense RNAs (1.8 % of non sex specific, 4 % of male specific and 2 % of female specific) or miRNAs (0% of non sex specific, 1 % of male and 0.4% of female) (Fig. 6B). If we filter out those genes that belong to these categories, the male specific gene list is enriched in terms associated to bacterial defense response (GO: 0019731, 0050830, 0042742), unfolded protein (GO: 0006986, 0034620), embryonic development (GO: 0000578, 0060429) and cytoskeleton assembly (GO: 0007015, 0014866). In the filtered list of female specific genes, we see enrichment in terms corresponding to courtship (GO: 0007619, 0008049, 0060179), development (GO: 0007484, 0048747, 0035215, 0035282, etc.), BMP signaling (GO: 0030510) and apoptosis (GO: 2001233) (Fig. 6C).

Next, we compared those genes that are significantly altered after up-regulated *miR-979* to those genes that are significantly altered because of LSE. In this case we consider LSE genes to consist of those belonging to our previously defined biological stress hallmark as well as those genes that are specific to maternal LL. Out of 3089 genes that are altered after maternal LL, regardless of the sex of the progeny, there are 97 genes (3.1 %) that are also altered after *miR-979* upregulation in both males and females. Attending to the effects in only males, out of 4681 genes affected in F1 males of F0 LL mothers, 408 genes (8.7 %) are also altered when *miR-979* is up-regulated in males. In contrast, of 5175 genes that are altered in F1 females of F0 LL mothers, 341 genes (6.6 %) show different expression in females with down-regulated *miR-979* (Fig. 6D).

In these gene lists, there is again a significant percentage of genes of undefined function or functional RNA transcripts (totaling 57 % in the non-sex-specific genes, 57 % in the male specific genes, 62 % in the female specific genes). In this case, if we filter out those terms, within the non sex-specific genes that are altered both in LSE conditions and with up-regulation of *miR-979* there is enrichment in GO terms associated to cellular response to UV, heat and light (GO: 0034644, 0034605, 0009411, 0071482) along with response to oxidative stress (GO: 0034599, 0032268). In the male specific list enriched GO terms correspond to antibacterial response (GO: 0019731, 0042742, 0050830) and oxidative stress response (GO: 0042743, 0034599) whereas in females enriched GO terms match glucose metabolism (GO: 0032024, 0042593, 0033500), courtship (GO: 0008049, 0007619), sensory perception (GO: 0007600, 0007606, 0050982, 0007608, 0050911) and ion transport (GO: 0006814, 0030001, 0015672). (Fig. 6E)

The presence of shared alterations after maternal LL conditions and up-regulation of *miR-979* further indicate that this could be one of the mechanisms by which maternal LL is inherited by the progeny. In addition, the role of *miR-979* regulation in altering processes that include oxidative stress response, humoral response and glucose metabolism, explains the differences in GB phenotypes that were previously described, further signaling *miR-979* as a risk factor for GB lethality, at least in part.

## Discussion

Here we have studied the effect of stress as a harmful lifestyle that impacts the *Brain fitness* of the next generation. Our results support the idea that the effects of maternal environmental stressors are inheritable to the progeny, as exemplified by glioblastoma development. GB grows faster and shows worse prognosis in affected animals. Notably, those inheritable changes in F1 brains depend on the stress type and the sex of the progeny. We have studied the effect of permanent light exposure as stressor, although other stressor modalities, including diet, are also reviewed. Using a *Drosophila melanogaster* experimental model of GB malignancy^8,112^, we observed that maternal luminic stress sensitizes F1 brains to GB progression. Permanent light exposure increases oxidative stress in the F0 germ line as well as in F1 brains, although the latter does not seem to have detrimental effect on F1 CNS development or F1 survival. By contrast, maternal stress reduces F1 lifespan and it accelerates GB progression including tumor invasion, proliferation and neurodegeneration. While stress impacts the *Brain fitness* of both sexes in the F1, RNAseq results suggest that the responses to inheritable changes have sex-specific and stressor-specific signatures. In this context, miRNAs emerge as a potential mechanism mediating this intergenerational effect, including *miRNA-979*.

## Stressors for *Brain fitness*

Prolonged light exposure, or light pollution, is part of daily lives and it can lead to behavioral and health disruptions, such as increased cancer risk^113–123^. In this work, we aimed to replicate normal indoor lighting (500-1,000 lux), to mimic a standard lifestyle condition of light exposure to inquire about the potential harmful effects. Thus, we optimized a permanent light protocol based on the exposure of flies to 24h of 1,000 luxes of white LED light for 7 days. In addition, we limited variables such as food, temperature, age of the fly, humidity, or amount of flies/tube. However, in nature this scenario is more complex with the sum of the simultaneous interaction of multiple sources of stress.

Light is the most important cue to synchronize the circadian clock and even levels of light at night as low as 4-5 lux of light during the dark phase or 0.2 lux under the door in a dark room) is sufficient to alter circadian rhythms in rats^124^ and also in flies^39,40^ For example, 3 days of dim light of 10 lux of light exposure is sufficient to promote circadian rhythm disruptions, neurodegeneration and worse prognosis in the Alzheimer’s disease (AD) model using *Drosophila melanogaster*^40^. In our work, we did not observe significant changes in circadian genes such as *Per* nor *Clock* in F1 females bearing maternal LL although we did observe increased expression in *Tim* and *Achl* genes (2.2 and 2.8-fold up-regulation respectively). In the case of F1 males, we observed no significant changes in those circadian genes. However, in both F1 females and males we found an increase in the expression of *Cry* (2.8 and 4-fold up-regulation respectively), a cryptochrome encoding a blue light photoreceptor that mediates light input^125^ and whose expression has been linked with GB progression^14^. Other neurodegenerative diseases conditions have been studied, such as Huntington’s disease (HD). HD mice models that were exposed to 20 lux light at night show a worse HD clinical progression^126^. For example, rats exposed to bright light (3,000 lux) for 20 days or 90 days showed a progressive reduction of tyrosine hydroxylase (TH)-positive neurons in the substantia nigra^127^. The authors suggest that this reduction could be triggered by dopamine oxidation due to the light cue that increases the presence of reactive oxygen species. Our results show that both F1 females and males show an increased expression of the gen *ple* (2.4 and 4-fold up-regulation respectively) a tyrosine 3-monooxygenase involved in DA synthesis, and also in both sexes of the gene *aaNAT1* (2.4 and 3.2-fold up-regulation respectively). The *aaNAT1* gene encodes an aralkylamine N-acetyltransferase involved in DA acetylation and oxidation in flies^128^. That is, the DA metabolism is increased in F1 bearing maternal LL and it might be related to the observed increase in brain oxidative stress^129^. A functional link may be that circadian rhythm disruption can be related to food intake^130,131^, which increases body weight in mice^131^ leading to a maternal risk factor for obesity and increased oxidative stress in oocytes^132^. Furthermore, additional hypotheses emerged including changes in the microbiota or hormonal status by LL ^133–138^. Thus, we propose that the effects of stress modulating *Brain fitness* are the result of a complex interaction between the inherited genetic preconditioning and GB development, so future experiments will likely shed light on the contributions to stress inheritance.

Epidemiological evidence determines that maternal suboptimal pre-conceptional or intrauterine exposure to smoking, stress, age, infection, chemicals and under- or overnutrition, impact offspring’s risk of, mainly metabolic, diseases throughout life^105^. Remarkably, in this work we used virgin females collected 0-3 days post-eclosion to abolish maternal age-dependent effects on the offspring and the potential stressful effect of conception to females^139^.

The current protocol used in this work is based on maternal stress, despite paternal health also playing a central role in this field of study^78,79^. Recent generational experiments in *Drosophila* showed that environmental impact on offspring might be parental sex-dependent^140^. Consistent with our hypothesis, maternal LL causes mitochondrial oxidative stress in the pre-conceptional female germ line and in the brains of the offspring. Maternal LL does not alter levels of expression of *Sod1* or *Cat* in F1 brains, two enzymes for antioxidant defense against ROS, suggesting that the pre-conditioned brain does not suffer oxidative damage. That could be explained by a circadian rhythmicity of antioxidant enzymes activity^141^ and it correlates with the normal CNS development in the progeny and survival results under physiological conditions except in the female context. It is worth noting that RNAseq results indicate enrichment in genes associated to oxidative stress in both male and female progeny, highlighting the need for further exploration of this mechanism in response to LL within the CNS. Regardless, LL seems to cause an effect on the ovaries, which could be a mechanism through which the stress is transmitted to the next generation, even if those animals are not directly exposed to the stressor. We suggest that the inheritance of this preconditioned brain (LSE) remains a stress hallmark or risk factor of society’s lifestyle that impacts health and pathology. Future experiments will be critical to determine if this inheritable risk for GB progression persists under chronic stress through generations as a transmissive maternal effect to have a more realistic model of modern society’s lifestyle.

## Genes for Brain fitness

How this LSE provokes stress- and sex-specific inheritable changes in F1 brains that accelerate GB growth and progression was the main question addressed in this work. The literature reports that the human male fetus is more susceptible to suboptimal *in utero* environmental exposures than females^142^. However, that depends on the stressor type and the duration of exposure, as well as on the features evaluated^143–145^. By profiling the effect of LL exposure on gene expression in the progeny, changes in gene expression were larger in offspring females compared to males descending from LL mothers. These changes include a sex-specific genetic hallmark, that includes genes involved in stimulus detection and metabolism, immune defense, and non-coding RNAs such as lncRNAs. Recent literature links the role of maternal inheritance of lncRNAs as a conserved generational mechanism that alters gene expression in the offspring upon different stressors and impact on offspring health^90,142,144–146^. Notably, there is a common core of 1416 genes among females and males to both experimental conditions (LL and HFD) as a general response to stress, that we defined as a hallmark of stress memory. Those genes are related to mitochondrial activity and transport, sexual reproduction, cellular metabolism, and cytoskeleton dynamics. Consistent with our results, the literature describes the stress response as an evolutionarily conserved mechanism that supports a suitable adaptative response and a specific genetic program in a changing environment^98–101,147^.

Among the specific genetic elements activated under maternal LL, we observed differences in gene expression levels that can explain the brain’s susceptibility to glioblastoma progression. Tumoral cells require specific metabolic traits to infiltrate and invade the healthy tissue through the communication between GB cells and neurons^95–97,146^. However, if this communication is disrupted, tumoral growth is impaired^95–97,146^. In this scenario, pre-conceptional maternal stress has been associated with alterations in the offspring developing neural and glial function and viability^104,148,149^, so it can explain how a preconditioned brain responds to a GB outcome. F1 individuals show up-regulation of genes linked to metabolic processes that are important to cancer cell survival and growth, including over 4-fold upregulation of expression of glutathione metabolism genes (*GstE9, GstS1*, *GstE11)* compared to controls, 2-fold up-regulation of expression of glucose metabolism genes (*Zw, Taldo, Rpi),* and up to 4-fold up-regulation of genes linked to lipid metabolism (*adp, Lip2, Lip4*). Moreover, F1 offspring show up-regulation of genes linked to mitochondria metabolism such as *Aldh*, an aldehyde dehydrogenase whose homolog in humans (*Aldh2*) is highly expressed in high-grade glioblastoma tissues^150^.There is also up-regulation of genes associated to signaling pathways that are relevant for glia-neuron communication such as wingless (*wg*).^97^When looking at genes that are specifically altered in F1 males, it is notable to find 2-fold down-regulation of genes associated to learning (*rut, dnc, graf, mnb*) and increased expression associated to bacterial defense response (*PGRPs, Dpts, imd, spz*). In contrast, F1 females show specific changes linked to sensory responses (*pyx, pain, piezo*), cell-cell communication (upregulated expression of actin and cadherin coding genes) and response to ROS (*Gst, Jra, p38a*). Moreover, mitochondrial redox genes such as *succinate dehydrogenase-A* (*Sdha*) or *cnc*, the ortholog of *human nuclear factor erythroid-derived 2-like 2* (*NFE2L2, Nrf2*) are 2-fold more highly expressed in F1 female brains than controls, suggesting higher activity of mitochondrion in F1 female brains that can contribute to pathogenesis^151^. In addition, female F1 brains report activation of genes involved in neuronal dynamics such as axon guidance, vesicular transport through MTs, synaptic activity or even neuronal differentiation, maturation and components related to neurogenesis. Overall, these findings support the idea that offspring have a different response to a sub-optimal pre-conceptional exposure and contribute to the risk of health problems.

In line with these results, we found genes that had been previously linked with glioma progression and glia-neuron communication to be enriched in F1 females from LL-F0 stressed females^96,97,146^. For example, genes associated to JNK signaling (*bsk, Tak1, kay, wnd, slpr)* are required for expansion of tumor microtubes in GB^97^and are upregulated in F1 females. Moreover, since GB flies show a disruption of circadian rhythms due to neurodegeneration^129^, we observed that circadian genes such as *Tim* or *Achl* show higher expression in F1 females. Another gene changes involve presynaptic genes such as *bruchpilot* (*Brp*), *liprin alpha* (*Lip-α*) and *synaptotagmin 1* (*Syt 1)* that participate in the establishment of glioma-glutamatergic neurons’ synapses as well as GB expansion and lethality, which show reduced expression compared to controls^146^. Moreover, genes related to aging are also upregulated in F1 females from LL-F0 stressed females such as *autophagy-related 1* (*Atg1*), *forkhead box, subgroup O* (*foxo*), *ribosomal protein S6 kinase* (*S6k*), and *mTOR* (*Tor*). Therefore, it is worth further evaluate if these genes increase the predisposition of the F1 female brain to develop an aggressive and lethal tumor by disrupting neuronal-GB and GB-GB cell communication and *Brain fitness*.

Lastly, our work and others in the field suggest the potential role of ncRNAs as a mechanism of generational inheritance^78,152–154^Additionally, ncRNAs respond to environmental exposures and therefore, are proposed as predictive biomarkers for specific stress conditions and disease progression^155^. Among these factors, as mentioned above, RNAs have potential value as downstream regulators of cellular responses through their levels of expression and, thereby, their targets’ expression. In this study, we observed that *miR-979* acts as a risk factor that impacts GB lethality and individuals’ survival. The down-regulation of *miR-979* prevents GB premature lethality in F1 flies bearing LSE. However, its upregulation has no impact on the GB phenotype. When comparing the effects of *miR-979* up-regulation on otherwise healthy brains, we are also able to recapitulate several of the genetic effects of LSE. This includes changes in expression of immune response genes in F1 males or sensory perception of stimuli and insulin response in F1 females. Although *miR-979* up-regulation alone is not capable of affecting all the pathways that are affected by LSE in GB progression, these commonly modified genes further validate the role of ncRNAs in transmitting stress memory across generations. This fact opens the possibility of personalized medicine and diagnosis, not only depending on the disease profile, but also on the inherited lifestyle or environmental conditions. If we could decipher this code and link it to favorable or detrimental conditions for *Brain fitness*, we will be able to trace the potential prognosis of a pathophysiological process in an organism, as an anticipated and personalized life prediction^156^. In addition, the potential use of RNA vaccines open new perspectives on modern medicine to modulate *Brain fitness* and disease response^157^.

Our findings suggest that female condition is a basal precondition for maternal LL effect. Shen et al., described that visible light had a strong sex-specific effect on mortality parameters in females versus males^158^, and that could also explain the differences in inheritance of light damage via maternal and not paternal lines in our data. Thus, biological sex remains an important factor for disease development. Moreover, although our results and others have demonstrated the generational inheritance of parental environmental conditions or lifestyle, it can also be asked if this pre-conceptional and inheritable state (preconditioned brain) is dynamic and susceptible to change through the life of an organism, despite its previous exposures. Therefore, *Brain fitness* might suggest the ability to use modifiable lifestyle habits as an intervention to reverse past experiences^159,160^. For example, Shen et al, observed that a high-protein content diet prevents the visible light toxicity mortality parameters in flies^159^. All these data together reinforce the urgent need to study modern lifestyle environmental factors as contributors to health and disease with an inclusive research perspective for better therapy design.

## Materials and methods

### *Drosophila* genetics, stocks, and maintenance

Drosophila strains were maintained at 25°C (unless specification) in standard food (SD) vials (unless specification) at constant humidity (60%) under a 12 hour-light/12hour-dark cycle (LD). Standard medium contained 5% yeast, 7,5% glucose, 1,125% agar, 5% wheat flour and 0,5% propionic acid.

### Acute Luminic Stress protocol (LL)

To test the inter/trans-generational effects of luminic stress (LL), 0-3 days-old virgin females were collected and exposed to different light regimes: (1) continuous white led-light (1000 luxes, LL) or (2) standard 24h regime of white led-light (12h light/12h darkness, LD) as control condition, for 7 days in standard food (SD) vials at 25° C (10 flies per tube). Flies were transferred into new SD vials every 2-3 days to avoid sticking in the food. To prevent the flies from getting stuck in the food, the vials were kept in horizontal position. At the end of the 7 days, female flies were returned to LD and were mated with young control males (F0) that were raised under LD and SD conditions at 25° C. Bottles contained 20 females and 20 males were placed in a 25° C-incubator under LD for 5 days to create de F1 generation. Flies from every subsequent generation were collected for survival experiments or kept on dry ice and storage at -20° C until RNA extraction for RT-qPCR and RNAseq experiments.

### Glioblastoma and LL protocol

We collected virgin females that carry the *UAS.PI3K ^CAAX^* and *UAS.EGFR ^λ^* and control virgin females that express *UAS.LacZ* and followed the above LL protocol for F0 females LL. Bottles contained 20 females and 20 males were placed in a 17° C incubator under LD conditions to create de F1 generation. After F1 flies eclosion, 0-3 days flies were transferred into new SD tubes and placed for 5 days or 7 days to develop a GB under LD regime at 29° C leading into inactivation of Gal80^TS^, allowing *Gal4* expression and tumor development. Then, flies were collected and dissected to evaluate the effect of maternal LL in the GB progression. In addition, some flies were collected on dry ice and storage at -20° C until RNA extraction for RT-qPCR experiments.

### High fat diet protocol (HFD)

To test the inter/trans-generational effects of high fat diet (HFD), *Drosophila* stocks were raised under different feeding regimes at 25° C under LD conditions: (1) high fat diet food vials containing standard food supplemented with 20% (volume) of coconut oil (HFD) or (2) standard diet (SD) as control condition. This protocol is optimized for *Drosophila* using a standard diet enriched in a 20% of coconut oil ^160^. To prevent the flies from getting stuck in the oily food, the vials were kept in horizontal position and a piece of paper was placed to the wall of the vial. After flies eclosion, 0-3 days old virgin females of both conditions were collected and mated with young control males (F0) that were raised under a SD feeding regime at 25 ° C in and LD conditions. Bottles contained 20 females and 20 males were placed in a 25° C-incubator at LD regime for 5 days to create de F1 generation. Flies from F1 generation were collected on dry ice and storage at -20° C until RNA extraction for RT-qPCR and RNAseq experiments.

### Survival assay

To evaluate the effect of maternal luminic exposure on offspring, we performed a survival assay with F1 males and females under control conditions and developing a GB. To do that, 1-7 days of F1 males and females of every luminic regime and experimental conditions were collected per genotype at 17°. Flies were placed into SD vials in groups of 8-10 flies and were kept at LD at 29° at constant humidity (60%). Flies were transferred into new standard food vials every 2-3 days to avoid sticking in the food. To prevent the flies from getting stuck in the food, the vials were kept in horizontal position.

### Mitotimer tool

0-3 day old adult female *Drosophila* were exposed to LL. Then, we collected the adult flies and the germ line was dissected and fixed for 20 minutes in 4% FA. Then, samples were washed 3 times for 10 minutes and we mounted the ovaries in Vectashield mounting medium with DAPI (Vector Laboratories, Inc.) and processed in the confocal microscopy. It is likely that fluorescence signal no longer lasts more than a few hours, thus implies an immediate imaging process after dissection. Moreover, we used this tool to measure intergenerational oxidative stress inheritance in the brain of F1 flies from 0-3 days-old adult female flies exposed to LL.

### RNA extraction, reverse transcription

10–15 adult head tissue samples were treated and homogenized with Trizol (Ambiend for Life techonologies). Chloroform was added and then centrifuged 13000 rpm at 4°C for 15 minutes. After discarding the supernatant, the RNA was treated overnight at -20°C with Isopropanol and then centrifuged 13000 rpm at 4°C for 10 minutes and washed with 75% Ethanol. RNA pellet was dissolved in DNAase RNAase free water and total RNA concentration is measured by using NanoDrop ND-1000.

### Microarray, RNAseq and bioinformatic analyses

We performed the LL protocol and collected the F1 samples. We extracted mRNA from the brains of progeny coming from females exposed to continuous light. Microarray results showed changes in mRNA levels of at least 39 genes (p<0.05 done in triplicates). This analysis was done by the company Flychip (Cambridge, UK). https://www.sysbiol.cam.ac.uk/CSBC/FlyChip.

Initial processing of RNAseq data was performed by the Omics Laboratory in the Cajal Institute under Dr. Jaime Pignatelli. This included initial quality control with FastQC (v 0.11.9) and MultiQC (v 1.13) and alignment to NCBI reference genome dm6 (https://www.ncbi.nlm.nih.gov/datahub/genome/GCF_000001215.4/) with HISAT2 (v 2.1.0). Transcripts were then quantified with HTSeq (v 0.11.2) software.

Having data for one individual per experimental combination, we next compared genes that showed changes in expression depending on the source of maternal stress and the sex of the progeny. Genes were considered to be sufficiently altered when gene expression changed to at least double or half the expression of the respective control (absolute logFC>1). These altered genelists for the different conditions (LL vs LD in males vs females; HFD vs SD in males vs females) were then compared for coinciding genes and represented in Venn diagrams. Enrichment analysis was then performed using a python wrapper of the Enrichr software (gseapy v1.1.8), to find GO terms that were significantly enriched in the different gene lists. When given, up or down-regulation of GO terms was approximated with the mean logFC values of all the genes that mapped to a GO term for a given comparison.

A second RNAseq was performed from samples of males and females with and without altered *miR-979* expression (repo> Cs*; repo> UAS. miR-979*). mRNA preparation was executed as before, while RNAseq library prep and sequencing were done by Macrogen Inc. Initial processing was also done at Macrogen, using Trimmomatic (v0.38) for removal of low quality reads, HISAT2 (v2.1.0) for alignment to reference genome *dm6* and feature counting with StringTie (v2.1.3b).

In this case, the presence of experimental condition duplicates allowed for identification of differentially expressed genes (DEGs) with EdgeR (v3.40.2). Library size factor bias was removed using the innate TMM method (*calcNormFactors*). Genes were considered significantly differentially expressed when p-values<0.05 were obtained in the Likelihood Ratio Test (*glmLRT*) with Sex as an interaction (*design <-∼Condition*Sex*), and additionally filtered by absolute logFC>1 compared to control conditions. These lists of DEGs were also studied for enrichment in particular GO terms with the python Enrichr wrapper (gseapy v1.1.8) as before and compared to gene lists resulting from the LL/LD contrasts.

### Immunostaining

To dissect larval brains, 3^rd^ instar larvae are collected and transferred into a dissection plate with phosphate buffered saline (PBS). Abdominal area is clamped with two dissection forceps and the genital area is removed. The tweezers are then inserted through the jaw, and with the help of the other forceps the body is rolled up until the tissue and internal organs are exposed **(Fig. M4)**. Then, the samples are transferred to an eppendorf with 4% formaldehyde (FA) to fix them for 20 minutes. After that, the FA is removed from the eppendorf and replaced by triton saline buffer 0.3% (PBS + 0,3% Triton) for the three-10 minutes wash at room temperature. Finally, samples are incubated on blocking (PBT 0.3% BSA 5%) for 30 min at room temperature snd incubated O/N at 4° C with primary antibodies: anti-repo (mouse) (1:200) that recognizes glial nuclei, anti-elav (mouse) (1:100) specific for neurons. After the three-15 minutes washing in PBT, samples are incubated in blocking buffer solution and secondary antibodies for two hours at room temperature. Finally, three-15 minutes washes are made with PBT, and brains are incubated O/N at 4° C with Vectashield mounting medium with DAPI (Vector Laboratories, Inc.). Then, samples are mounted in coverslips with Vectashield mounting medium with DAPI (Vector Laboratories, Inc.) and stored at 4° C.

To quantify the number of synapses After removing internal organs and adipose tissue, the PBS is removed, and the sample is fixed in 4% formaldehyde (FA) for 5 minutes at room temperature. FA is removed from the dissection plate and replaced by triton saline buffer 0.3% (PBS+ 0,3% Triton). Tungsten pins are carefully removed, and the larva is transferred to a 4-well plate, where three-10 minutes washes are done with PBT at room temperature and the incubation with blocking buffer solution (PBT+ BSA 5%) for 30 minutes at room temperature. Synapses are revealed when using α-Nc82 antibody (Bruchpilot) mouse in PBT O/N at 4° C. In addition, samples are stained with polyclonal antibody α-Hrp (Horseradish peroxidase) rabbit (1,300), labeling the neuronal membrane **(Fig. M5 C)**. After three-10 minutes washing with PBT 0.3%, samples are incubated for 2 hours at room temperature with the Alexa 488 mouse fluorophore (1: 500) for the detection of Bruchpilot, and the fluorophore Alexa 654 anti-rabbit. Then, samples are mounted in coverslips with Vectashield mounting medium (Vector Laboratories, Inc.) and stored at 4° C.

To report mitochondrial oxidative stress, we dissected adult *Drosophila* ovaries and brains. Samples are fixed in 4% formaldehyde (FA) for 20 minutes at room temperature in a 4-well plate. FA is removed and replaced by triton saline buffer 0.3% (PBS+ 0,3% Triton) for three-10 minutes washed at room temperature. Finally, we incubated the samples O/N with PBT and Vectashield mounting medium with DAPI (Vector Laboratories, Inc.) Then, samples are mounted with Vectashield mounting medium with DAPI and stored at 4° C.

### Image and statistical analyses

The images of *MitoTimer* reporter r were analyzed using the Plot Profile tool from FIJI (ImageJ 1.52v) software. This display allows to measure pixel intensity from green or red signal coming from ovaries and brains using equivalent position in the z axis of ovaries and brain lobes per sample.

To measure glial membrane, we used a *UAS-CD8-GFP* or *UAS-myristoylated-RFP* reporters under *repo-Gal4* expression control. Measured glial volume using Imaris surface tool (Imaris 6.3.1 software). To quantify glial cell numbers and synapses number, glial nuclei are stained with anti-repo antibody and active zones in the NMJ with Nc82 monoclonal antibody. The Imaris spots tool (Imaris 6.3.1 software) quantifies the number of repo positive cells and synapses number.

Statistics were analyzed using Graphpad Prism 8 software https://www.graphpad.com/. We performed the normality test using D’Agostino and Pearson omnibus normality test in order to perform the more suitable test for each experimental set. Student’s t-test and ANOVA test with Bonferroni’s post-hoc for parametric parameters. For non-parametric data, Mann-Whitney test and Kruskal-Wallis test with Dunns post-hoc are used. The survival assays are analyzed with Mantel-Cox test. Statistical analyses are included in each figure description. The null hypothesis were accepted when p values were *p<0,05; **p<0,01, ***p<0,001.

### Declaration of Generative AI usage

Declaration of generative AI and AI-assisted Technologies in the writing process: During the preparation of this work the author(s) used ChatGPT with GPT-4 to improve readability and language. After using this tool/service, the author(s) reviewed and edited the content as needed and take(s) full responsibility for the content of the publication.

## Acknowledgements

We thank Dr. Alberto Ferrús and Dr. Francisco A. Martín for their careful reading of the manuscript and insightful scientific discussions. We are grateful to the Advanced Microscopy Unit at ISCIII for technical support and to the Bloomington Drosophila Stock Center (BDSC) and Vienna Drosophila Resource Center (VDRC) for providing fly stocks. This work was supported by the project PID2022-139786OB-I00 Spanish Ministerio de Ciencia, Innovación y Universidades to SCT.

---

**Supplementary material 1.**
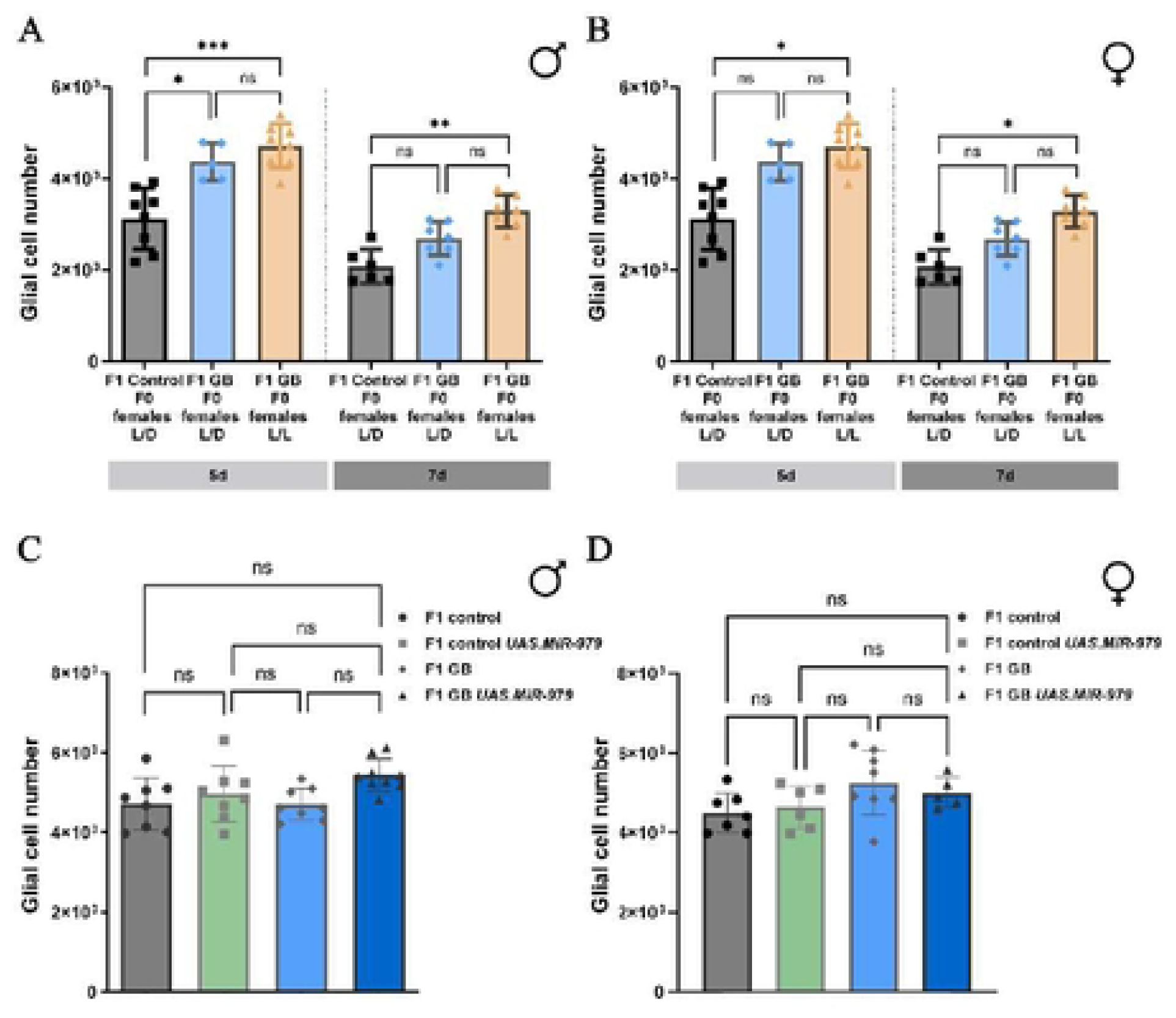
Quantification of glial cell number at 5 days and 7 days of tumor development in F1 males(A) andF1 females(B) One-way ANNOVA test, Bonferroni is multi pie comparition pos-test and One-way Kruskal-Wallis test Dunnis multiple comparition post-test * p<0,05., ** p<0,005, *** p<0,001.Quantification of glial cell number at 5 days of tumor development in F1 males(C) and F1 females (D) of F1 control (repo-Gal4> tubGal80TS CS), F1 control UAS.MiR-979 (repo-Gal4> tubGal80TS; UAS.MiR-979), F1 GB (repo-Gal4> tubGal80TS, UAS-dEGFRZ, UAS-dp110CAAX), F1 GB UAS.MiR-979 (repo-GalΦ> tubGal80TS, UAS-dEGFRZ, UAS-dp110CAAX; UAS.MiR-979). One-way ANNOVA tes, Bonferroni is multiple comparition pos-test

